# Habenular and striatal activity during performance feedback are differentially linked with state-like and trait-like aspects of tobacco use disorder

**DOI:** 10.1101/771915

**Authors:** Jessica S. Flannery, Michael C. Riedel, Ranjita Poudel, Angela R. Laird, Thomas J. Ross, Betty Jo Salmeron, Elliot A. Stein, Matthew T. Sutherland

## Abstract

Although tobacco use disorder is linked with functional alterations in the striatum, anterior cingulate cortex (ACC), and insula, preclinical evidence also implicates the habenula as a contributor to negative reinforcement mechanisms maintaining nicotine use. The habenula is a small and understudied epithalamic nucleus involved in reward and aversive processing that is hypothesized to be hyperactive during nicotine withdrawal thereby contributing to anhedonia. In a pharmacologic fMRI study involving administration of nicotine and varenicline, two relatively efficacious cessation aids, we utilized a positive and negative performance feedback task previously shown to differentially activate the striatum and habenula. By administering these nicotinic drugs (vs. placebos) to both overnight abstinent smokers (*n*=24) and nonsmokers (*n*=20), we delineated feedback-related functional alterations both as a function of a chronic smoking history (trait: smokers vs. nonsmokers) and as a function of drug administration (state: nicotine, varenicline). We observed that smokers showed less ventral striatal responsivity to positive feedback, an alteration not mitigated by drug administration, but rather correlated with higher trait-level addiction severity among smokers and elevated self-reported negative affect across all participants. Conversely, nicotine administration reduced habenula activity following both positive and negative feedback among abstinent smokers, but not nonsmokers; greater habenula activity correlated with elevated abstinence-induced, state-level tobacco craving among smokers and elevated social anhedonia across all participants. These outcomes highlight a dissociation between neurobiological processes linked with the trait of dependence severity and with the state of acute nicotine withdrawal. Interventions simultaneously targeting both aspects may improve currently poor cessation outcomes.

**One-sentence teaser:** In a pharmacological fMRI study, e dissociate brain alterations in the habenula linked with nicotine withdrawal and striatal alterations linked with addiction.

## INTRODUCTION

The likelihood of a cigarette smoker successfully quitting on a given quit attempt is low [1, 2], with numerous attempts often required to achieve cessation [3]. Poor treatment outcomes for tobacco use disorder testify to the addiction liability of nicotine which, like other drugs of abuse, leads to increased local concentration of dopamine (DA) within the mesocorticolimbic (MCL) circuit when acutely administered [4]. Nicotine stimulates midbrain DA neurons projecting primarily to striatal and prefrontal brain regions through agonist effects on nicotinic acetylcholine receptors (nAChRs) [5–7]. During the initiation of use, acute nicotine-induced DA release is thought to positively reinforce continued drug seeking and taking [8]. As initial use transitions to chronic smoking, neuroplastic alterations within the MCL circuit and beyond are thought to induce a condition of dependence/addiction accompanied by a shift towards negative reinforcement mechanisms perpetuating continued use [9, 10]. Abrupt smoking cessation perturbs homeostasis maintained in the presence of chronic nicotine, giving rise to the nicotine withdrawal syndrome. Hallmark features of the syndrome, serving as major barriers to short-term cessation, include mood disturbances, attentional & cognitive impairments, and reward processing alterations [11–14].

Reward processing alterations frequently observed among dependent smokers manifest as both striatal hypo-responsivity to nondrug rewards [15–22] and striatal hyper-responsivity to drug-related stimuli [23, 24]. These alterations are thought to lead to the prioritization of nicotine over other rewards [25] and greater alterations are linked with worse cessation outcomes [18, 21, 26, 27]. Nicotine delivery via smoking ameliorates acute abstinence-induced dysregulation of affective [28], cognitive [29], and reward processes [30], thereby perpetuating smoking via negative reinforcement [31]. However, emerging evidence suggests a distinction in the neural responses contributing to smoker’s dysregulated reward processing such that nicotine administration may normalize blunted striatal activity linked with *reward anticipation* [15, 26], but not with *reward receipt* [22]. As opposed to being modulated by acute nicotine administration, blunted striatal responsivity to reward receipt has been linked with measures of chronic nicotine exposure including scores on the Fagerström Test of Nicotine Dependence (FTND) [22] and years of daily smoking [16]. Accordingly, we have previously proposed that blunted striatal responsivity is linked with trait-level addiction severity whereas other neurobiological circuits (e.g., those centered on the insula) are linked with state-level nicotine withdrawal [32]. Distinguishing the neural underpinnings of tobacco use disorder that are, and are not, mitigated by acute nicotine administration may expedite development of improved smoking cessation interventions for withdrawal management and/or relapse prevention.

While the striatum, anterior cingulate cortex (ACC), and insula are regarded as key constituents in the neurocircuitry of addiction [8, 9], emerging preclinical evidence also implicates the habenula as a contributor to negative reinforcement mechanisms perpetuating nicotine use [33–36]. The habenula is a small epithalamic nucleus that integrates information from limbic forebrain regions to modulate midbrain structures involved in monoamine neurotransmission and thus has been linked with reward, anxiety, stress, cognitive, and motor processes [37]. The habenula is divided into lateral and medial parts. The lateral habenula plays an important role in reward processing, specifically, when an expected reward is omitted, increased activity in the lateral habenula is thought to inhibit DAergic midbrain cells leading to decreased DA signaling in the striatum [37, 38]. Moreover, the medial habenula is of particular interest in the context of nicotine addiction as it possesses a high density of nAChRs [39] and has been functionally linked with nicotine self-administration [40, 41] as well as the aversive effects accompanying acute nicotine withdrawal [42] and high nicotine doses [43]. At the resolution of fMRI studies, medial and lateral aspects cannot be dissociated and indeed distinguishing habenula signals from those of the surrounding thalamus is difficult. However, only the habenula responds to reward prediction, (non-)reward outcomes, and performance feedback [44–47]. For ease of presentation, below we use the term habenula while acknowledging that the fMRI signal from such a small anatomical region may be contaminated by non-habenula signals (e.g., other thalamic regions, physiologic noise).

The habenula is thought to play a critical role in the transition from positively reinforced, initial drug exposure to negatively reinforced, compulsive drug use and addiction [34, 48, 49]. According to this perspective, repeated exposure to addictive drugs, including nicotine, is accompanied by increasingly elevated lateral habenula activity and the recruitment of medial habenula activity which both contribute to the dysregulation of DAergic circuits [34, 35, 38]. While drug use may initially decrease lateral habenula activity thereby contributing to the positive reinforcement of drug-taking, as use continues, neuroadaptations occur potentially leading to elevated lateral habenula activity via allostatic processes [34]. Elevated lateral habenula activity may contribute to an acute hypodopaminergic withdrawal state and chronic drug use may then serve as a means to reduce this habenular hyperactivity [34, 35]. In addition, the medial habenula modulates nicotine-induced DA release and associated motivational processes in rodent models [50, 51]. Thus, elevated lateral habenula activity and the growing involvement of the medial habenula are thought to disproportionally prioritize nicotine rewards over other rewards [34]. Despite preclinical evidence and theoretical perspectives linking habenula function with the development and maintenance of nicotine addiction, to our knowledge, the impact of chronic and/or acute nAChR stimulation on habenula activity has not been characterized in humans.

While the habenula’s small size generally limits its assessment in human fMRI studies [52], we utilized a performance feedback task previously shown to differentially activate the habenula, ACC, insula, and ventral striatum following positive and negative performance feedback [45]. As current pharmacologic smoking cessation aids are only modestly effective, elucidating the impact of early nicotine withdrawal and pharmacotherapy administration on the activity of these brain regions may facilitate identification of neurobiological targets for improved interventions. Nicotine replacement therapy (NRT) and varenicline are two modestly efficacious and currently available pharmacologic cessation aids. Whereas nicotine is regarded as a full agonist at ⍰4β2 nAChRs, varenicline is regarded as a partial agonist at these receptors [53]. In the context of a pharmacologic-fMRI study, we collected neuroimaging data from biochemically-verified overnight abstinent smokers and nonsmokers (**Supplemental Table S1**) while they completed a performance feedback task following administration of nicotine, varenicline, neither, or both. This experimental design allowed us to dissociate functional brain alterations associated with a chronic smoking history (smokers vs. nonsmokers) from those associated with pharmacologic administration (nicotine, varenicline). We addressed three main empirical questions involving task-, group-, and drug effects. Regarding task effects, we expected brain activity patterns associated with the performance feedback task to replicate findings from the original implementation [45], specifically increased habenula, ACC, and insula activity following negative feedback and increased ventral striatal activity following positive feedback. Regarding group effects, we expected that smokers (versus nonsmokers) would display decreased striatal activity to positive feedback indicative of neuroplastic changes associated with chronic nicotine exposure. Finally, regarding drug effects, we expected that nicotine and varenicline administration would modify acute withdrawal-induced activity in the habenula and other regions associated with processing performance feedback (i.e., positive and/or negative outcomes) among smokers, but not nonsmokers (who were of course, not in the state of nicotine withdrawal).

To these ends, we collected self-report questionnaire, behavioral task performance, and MRI data from participants in the context of a within-subject, double-blind, placebo-controlled, crossover study, involving two drugs: transdermal nicotine (NicoDerm CQ, GlaxoSmithKline) and oral varenicline (Chantix, Pfizer). Overnight-abstinent smokers (∼14 hours) and nonsmokers both completed 6 fMRI visits on different days over a 6-8 week study period (Fig. 1A). At three time points during a varenicline administration regimen (PILL factor: pre-pill [baseline] vs. placebo vs. varenicline), all participants completed MRI scanning on two occasions, once while wearing a nicotine patch and once while wearing a placebo patch (PATCH factor). Participants completed self-report questionnaires to quantify clinically-relevant constructs, including trait-levels of addiction severity (FTND), state-levels of tobacco craving (Tobacco Craving Questionnaire), negative affect (Positive and Negative Affect Schedule), and social anhedonia (Revised Social Anhedonia Scale). To probe striatal and habenular function, we employed a performance feedback task [45] in which participants predicted which of two moving balls, starting from different locations and traveling at different speeds, would reach a finish line first after viewing a short sequence of the balls’ motion (Fig. 1B). Task difficulty was dynamically and individually adapted thereby maintaining error rates at ∼35% so that participants remained uncertain about their performance until feedback presentation. Participant button-press responses (correct vs. error) were followed by feedback that did or did not provide information about trial outcomes (informative vs. non-informative). Task behavioral performance measures included the percent of *correct*, *error*, and *no response* trials and response times. Neuroimaging task effects were assessed in a whole-brain dependent samples *t*-test (informative-correct vs. informative-error trials), group effects were assessed in an independent samples *t*-test (smokers vs. nonsmokers), and drug effects in a linear mixed-effects framework (GROUP * PATCH * PILL).

**Figure 1.**
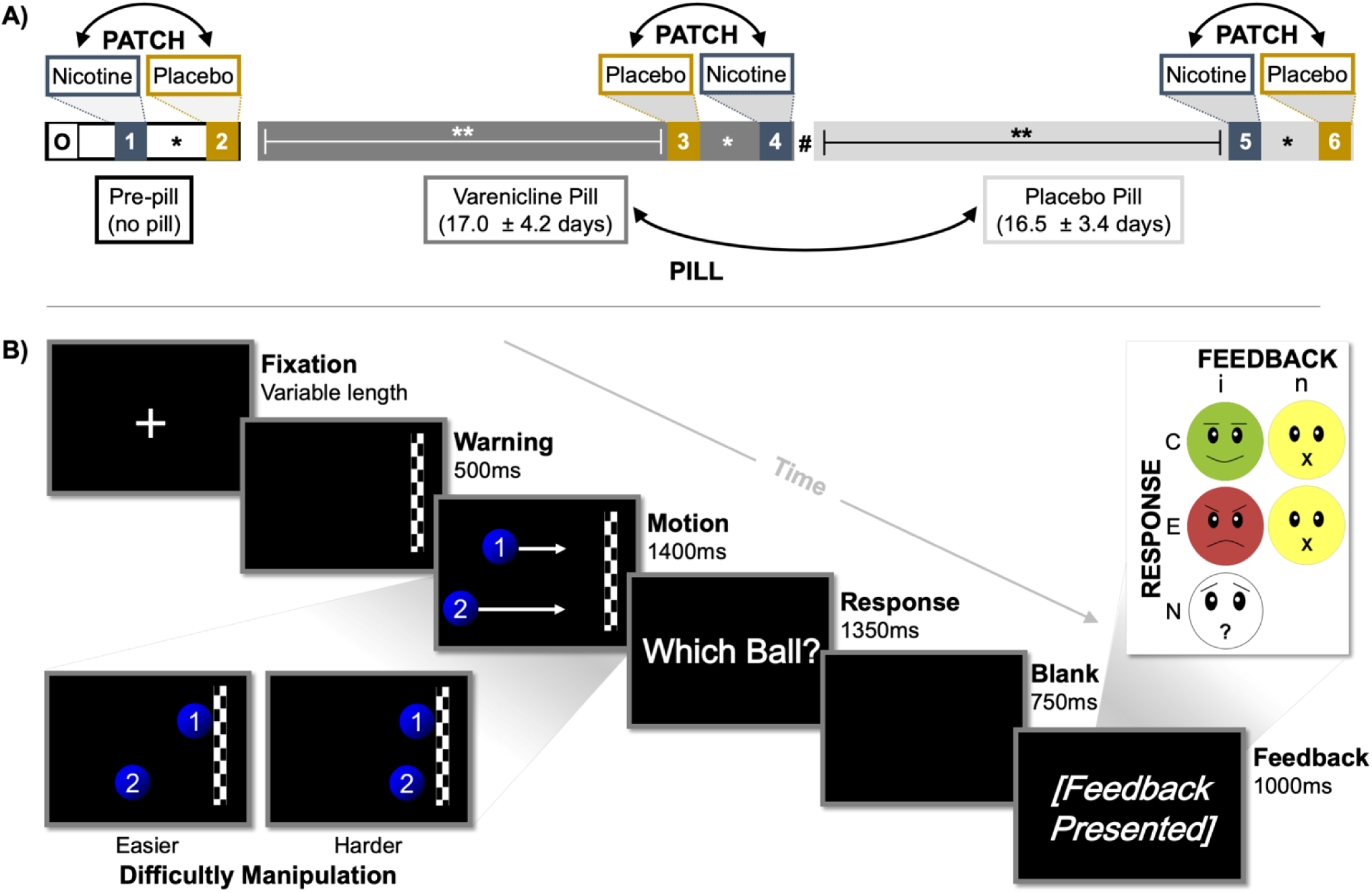
Study Overview Schematics. **<u>(A)</u>** Illustration of pharmacological study design. Following an orientation session (O), all participants completed 6 fMRI assessments over the course of the 6-8 week study duration. Before the onset of a study pill regimen (pre-pill), participants completed assessments wearing transdermal nicotine and placebo patches on different days. Subsequently, participants underwent varenicline (mean ± SD: 17.0 ± 4.2 days) and placebo pill administration (16.5 ± 3.4 days) and again completed nicotine and placebo patch scans towards the end of both PILL periods. Double-head arrows indicate the randomization and counterbalancing of drug orders across participants. * Nicotine and placebo patch scan sessions were separated by 2.9 ± 1.7 days. ** Neuroimaging assessments occurred 13.9 ± 2.3 days after the onset of each PILL period. Neurocognitive assessments were conducted one week after the onset of each PILL period. # A washout interval did not separate varenicline and placebo pill epochs. **<u>(B)</u>** Illustration of performance feedback task. Following a variable *fixation* interval and a *warning* stimulus indicating an upcoming event, participants viewed a sequence of two balls moving across the screen at different speeds from different starting locations. After a short ‘video’ clip of *motion*, the balls disappeared before reaching the finish line and participants indicated via a button press response *which ball* they believed would have reached the finish line first. Performance *feedback* was delivered at the end of each trial in a two-factor FEEDBACK (informative [i] vs. non-informative [n]) * RESPONSE (correct [C] vs. error [E]) fashion. A fifth feedback type was presented when participants failed to give a response (*no response* [N] trials). Task difficulty was dynamically manipulated on each trial and for each participant by modifying the time difference between the two balls’ intended arrival at the finish line.

## RESULTS

### Behavioral Measures

#### Task Effects

Confirming that task difficulty was dynamically and individually tailored, participants responded correctly on 60.7 ± 0.7% (mean ± SEM), erroneously on 35.5 ± 0.2%, and failed to respond on 3.8 ± 0.6% of trials (Fig. 2A). Response times were significantly longer for *error* (607 ± 14ms) versus *correct* trials (570 ± 13ms, *F*[*1, 43*] = 130.4, *p* < 0.001, Fig. 2B) across both informative and noninformative feedback conditions indicating a higher degree of uncertainty on error trials. These outcomes are consistent with the original task implementation [45] and the interpretation that the dynamic difficulty manipulation rendered participants dependent on feedback (as opposed to self-monitoring) for evaluation of their trial performance.

**Figure 2.**
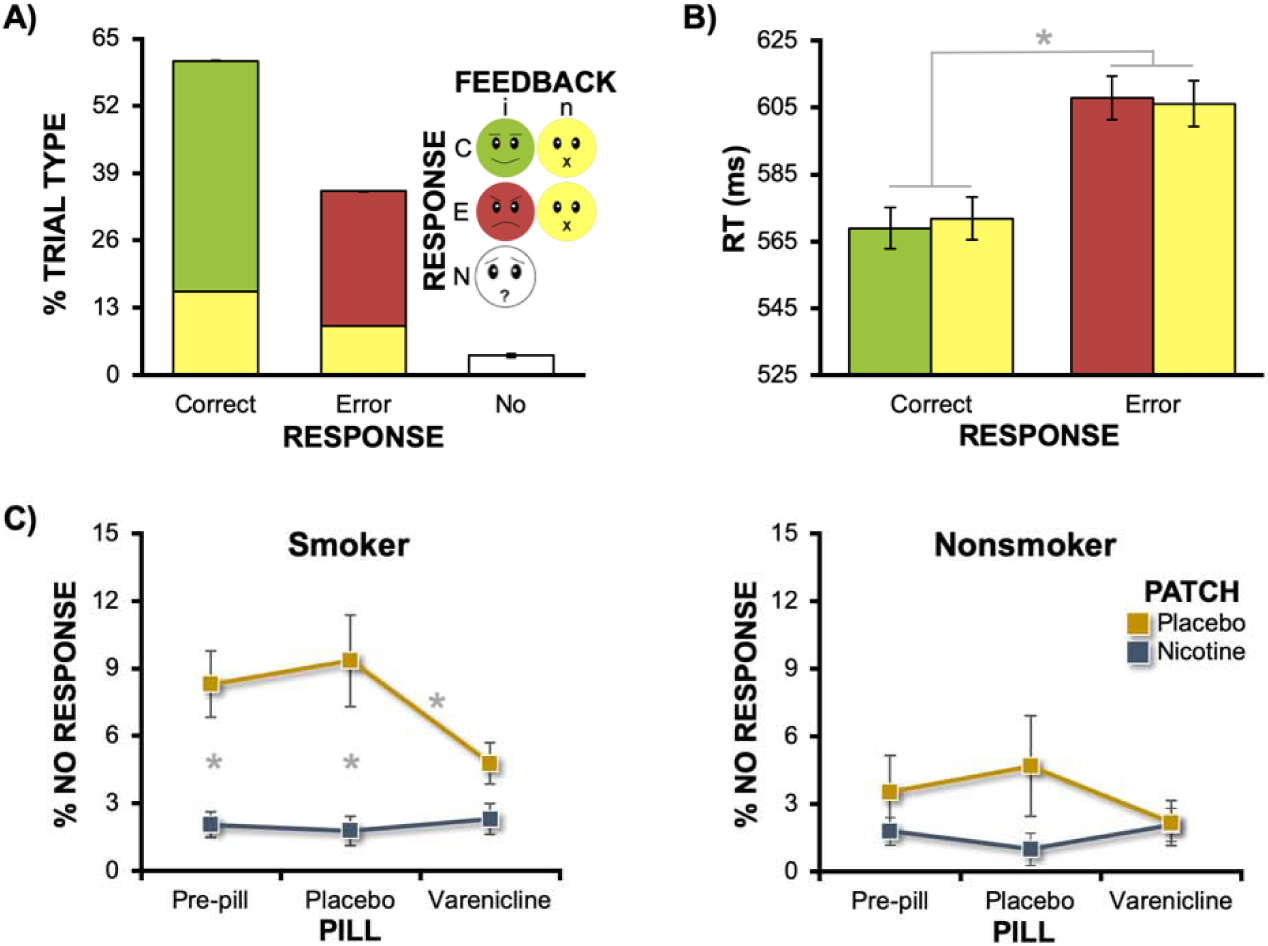
Task behavioral metrics as a function of trial type, group, and drug conditions. <u>(**A**)</u> Task difficulty was individually tailored to achieve consistent percentages of *error* trials across participants and sessions. Following both error and correct responses, informative feedback was presented on 73% (red/green) and noninformative feedback on 27% of trials (yellow) receiving a participant response. The *no response* trials (white) reflected momentary lapses of attention and was not under the control of the task’s adaptive algorithm. <u>(**B**)</u> Average response time (RT) for errors was longer than that for correct trials (RESPONSE main effect [* *p* < 0.001]). <u>(**C**)</u> Nicotine and varenicline decreased the percent of *no response* trials among both smokers (PATCH * PILL: *p* = 0.02) and nonsmokers (PATCH * PILL: *p* = 0.049). * = Significant *post hoc* pairwise comparisons at a Bonferroni-corrected threshold (*n* = 3 comparisons). Error bars = SEM.

#### Group and Drug Effects

We assessed the percent of *no response* trials, which we conceptualized as a gross behavioral measure of attention, in a GROUP * PATCH * PILL mixed-effects ANOVA (Fig. 2C). Regarding group effects, we observed modest, but non-significant smoker versus nonsmoker differences when considering the percent of *no response* trials (smokers: 4.8 ± 0.8%; nonsmokers: 2.5 ± 0.9%; *F*[1, 42] = 3.5, *p* = 0.07). However, this GROUP main effect was qualified by the presence of a significant GROUP * PATCH interaction (*F*[1, 42] = 5.6, *p* = 0.02) which was then followed by separate within-group, repeated-measures ANOVAs with PATCH and PILL as factors. Among abstinent smokers, both nicotine and varenicline impacted performance as indicated by a significant PATCH * PILL interaction (*F*[1.7, 38.8] = 4.6, *p* = 0.02, Fig 2C). Specifically, nicotine-induced decreases in *no response* trials were observed in the absence of varenicline under both the pre-pill (*t*[23] = −3.5, *p* = 0.006) and placebo-pill conditions (*t*[*23*] = −4.0, *p* = 0.003). Similarly, a varenicline-induced decrease was also observed (in the absence of nicotine under the placebo-patch conditions) when comparing active varenicline-pill versus placebo-pill conditions (*t*[*23*]= −2.8, *p* = 0.03). Among nonsmokers, similar, albeit less robust effects of nicotine and varenicline were observed as indicated by a significant PATCH * PILL interaction (*F*[1.7, 32.6] = 3.6, *p* = 0.049, Fig. 2C), although no *post hoc* pairwise comparisons reached significance at a Bonferroni-corrected threshold. Taken together, these outcomes provided confirmation that a) nicotine and varenicline administration were linked with alterations in task behavioral responding among both cohorts, and b) both drugs had their greatest effects among abstinent smokers, thereby serving as a design and pharmacologic-manipulation check.

### Imaging Measures

#### Task Effects

To characterize brain activity differentially modulated by performance feedback, we contrasted whole-brain BOLD signal changes associated with informative (negative) feedback following errors (iE) versus informative (positive) feedback following correct (iC) trials. Across all participants and sessions, negative feedback yielded greater activity (iE > iC) notably in the thalamic region encompassing the habenula, bilateral anterior insula, and the ACC extending into pre-supplementary motor area (pre-SMA) and SMA (Fig. 3A, **Supplemental Table S2**; see **Supplemental Fig. S1** for additional details on the anatomical definition of the habenula). Conversely, positive feedback yielded greater activity (iE < iC) in the bilateral ventral striatum (nucleus accumbens). Isolating the critical task manipulation to the type of feedback (i.e., negative vs. positive) as opposed to the type of response (i.e., error vs. correct trials), BOLD signal change was similar between error and correct trials followed by noninformative feedback (nE = nC) (Fig. 3B, **Supplemental Fig. S2**).

**Figure 3.**
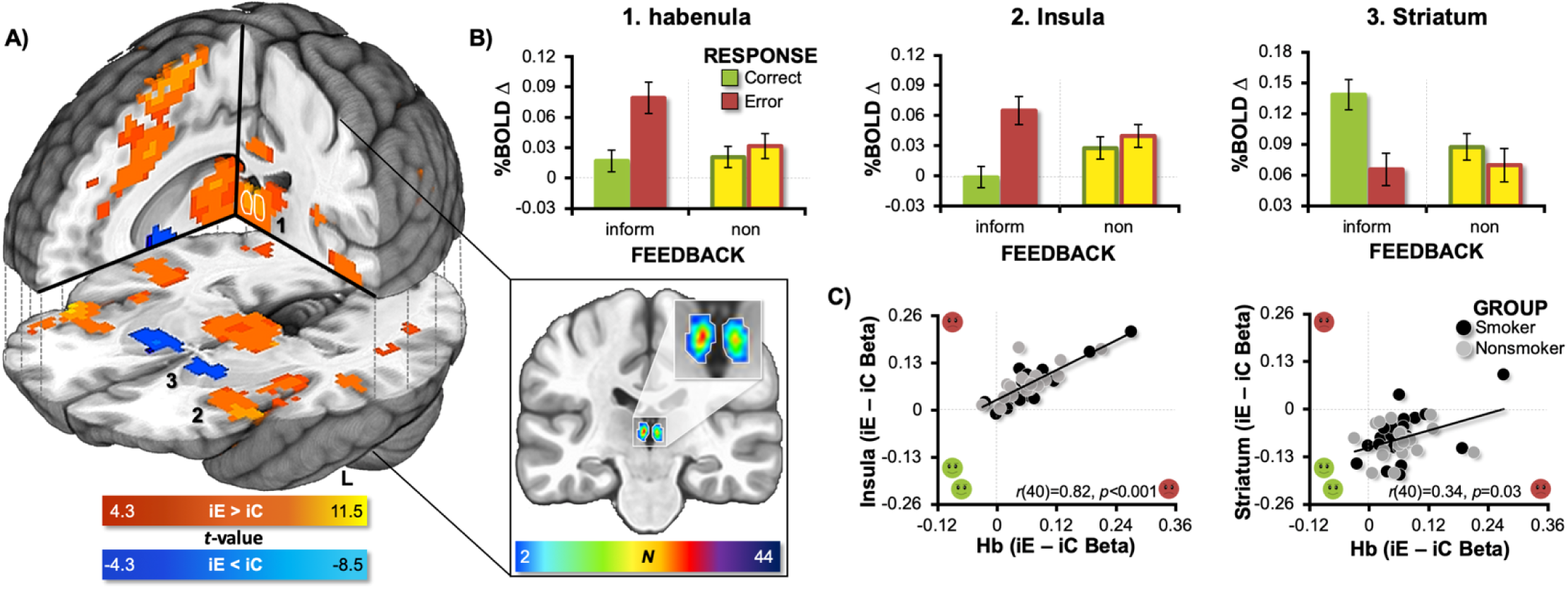
Task effect: Whole-brain activity following negative (iE) and positive (iC) feedback. <u>(**A**)</u> Negative feedback increased activity notably in the thalamus encompassing the habenula (outlined in white), bilateral insula, and ACC extending into the pre-SMA and SMA (warm colors = iE > iC feedback, *p_corrected_* < 0.001). Positive feedback increased activity in the bilateral ventral striatum (cold colors = iE < iC feedback). To provide an anatomical frame of reference (insert), we manually defined the habenular complex within each participant’s T1-weighted structural image utilizing anatomical landmarks and visible tissue contrast (detailed in **Supplemental Fig. S1**). These individual participant habenula masks were then normalized and summed to produce an overlap image with voxel values possibly ranging from 0 to 44 (24 smokers, 20 nonsmokers). We created an inclusive anatomical frame of reference by outlining those voxels with at least a 2-participant overlap (white outline). <u>(**B**)</u> Mean percent BOLD signal change (*β*) values from all four feedback conditions (iE, iC, nE, and nC trials) for the 1. habenula, 2. left anterior insula, and 3. left ventral striatum. While a selective statistical test of the RESPONSE * FEEDBACK interaction would constitute a circular analysis [60], see **Supplemental Fig. S2** for a whole-brain interaction test. <u>(**C**)</u> Correlations between the contrast values (iE – iC betas) from pairs of ROIs indicated that higher negative feedback responsivity in the habenula region (larger positive values) correlated with higher negative feedback responsivity (larger positive values) in the left insula (*p* < 0.001), right insula (*r*[40] = 0.85, *p* < 0.001, data not shown), and ACC/pre-SMA/SMA (*r*[40] = 0.90, *p* < 0.001, data not shown). Conversely, higher negative feedback responsivity in the habenula ROI (larger positive values), correlated with lower positive feedback responsivity in both the left striatum (smaller negative values, *p* = 0.03) and right striatum (*r*[40] = 0.31, *p* = 0.046, data not shown). See **Supplemental Table S2** for cluster coordinates.

Given the habenula’s known association with error processing circuits [45, 54, 55] and reducing dopaminergic activity [38, 46, 56], we explored for differential relationships between the feedback-related responsivity (iE – iC) of the habenula-containing ROI and the responsivity of other brain regions to either negative feedback (iE – iC > 0) or positive feedback (iE – iC < 0). Across all participants and sessions, as the negative feedback responsivity of the habenula increased, the insula’s negative feedback responsivity also increased (*r*[40] = 0.82, *p* < 0.0001). In other words, the more responsive the habenula was to negative feedback, the more responsive the insula also was to negative feedback. Conversely, as the negative feedback responsivity of the habenula increased, the striatum’s positive feedback responsivity decreased (*r*[*40*] = 0.34, *p* = 0.03) (Fig. 3C). In other words, the more responsive the habenula was to negative feedback, the less responsive the striatum was to positive feedback. Collectively, these task-effect results are consistent with contemporary views of the habenula’s central role in negative outcome processing and modulation of dopaminergic circuitry [57–59].

#### Group Effects

To elucidate alterations in brain activity linked with *chronic* smoking, we compared session-averaged [iE – iC] contrast images between smokers and nonsmokers within a composite mask of interest (**Supplemental Fig. S3**). Smokers showed <u>less positive</u> feedback responsivity in the bilateral ventral striatum yet <u>more negative</u> feedback responsivity in the left insula (Fig. 4A, B, **Supplemental Table S3**). Assessing beta weights separately for iE and iC trials indicated that smokers’ BOLD signal change following iC (positive) feedback was reduced by 60% in the left and 49% in the right ventral striatum, relative to that observed among nonsmokers (**Supplemental Fig. S4**). When assessing state-level drug effects in these clusters, we observed a high degree of consistency (i.e., non-significant differences) in the [iE – iC] contrast values across drug conditions and hence no indication of nicotine or varenicline effects (Supplemental Fig. S4). However, consistent with previous observations [22] and an effect of chronic nicotine dependence, larger reductions in ventral striatal responsivity to positive feedback among smokers correlated with higher FTND scores, a trait-level measure of addiction severity (Fig. 4C; left: *r*[21] = 0.52, *p* = 0.036; right: *r*[21] = 0.43, *p* = 0.13). Additional exploratory analyses across all participants suggested that reduced ventral striatal responsivity to positive feedback also correlated with higher session-averaged (i.e., trait-level) negative affect (left: *r*[40] = 0.34, *p* = 0.03; right: *r*[40] = 0.34, *p* = 0.03; Supplemental Fig. S4).

**Figure 4.**
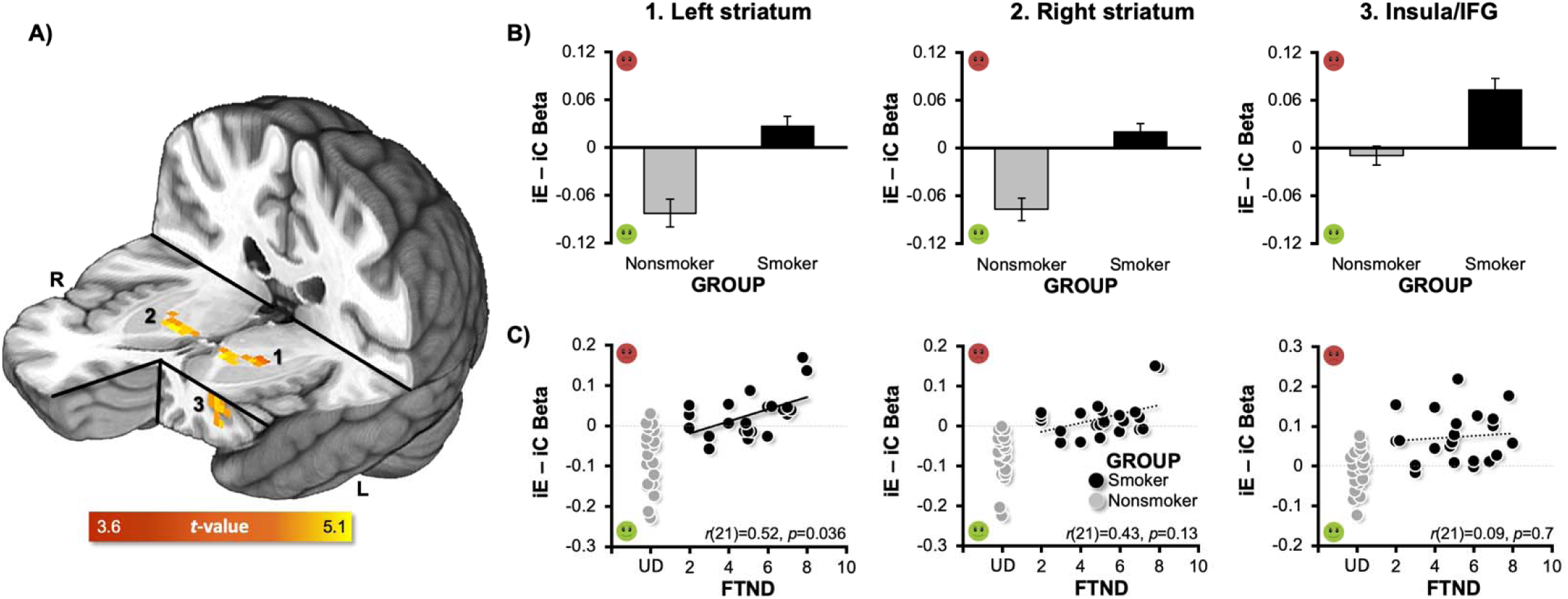
Group effect: Smoker versus nonsmoker differences in feedback responsivity. <u>(**A**)</u> Group differences in [iE – iC] contrast values were observed in the bilateral ventral striatum and left anterior insula/IFG (independent-samples *t*-test of session-averaged contrast values, *p_corrected_* < 0.05). <u>(**B**)</u> Chronic smoking was associated with reduced positive feedback responsivity (smaller negative/larger positive values) in the 1. left and 2. right striatum and increased negative feedback responsivity (larger positive values) in the 3. left insula/IFG. <u>(**C**)</u> Contrast values in smokers’ left ventral striatum correlated with Fagerström Test for Nicotine Dependence (FTND) scores (*p*_Bonferroni-corrected_ = 0.036), such that higher levels of addiction severity were linked with greater alterations in positive feedback responsivity (black circles = smokers, gray circles = nonsmokers). Indicative of regional specificity, a similar correlation with FTND scores was not observed for the left anterior insula/IFG (*p* = 0.7). All nonsmoker FTND scores are undefined (UD) and are staggered around that point in the graph to allow for visualization of all data points. See **Supplemental Table S3** for cluster coordinates and **Supplemental Fig. S4** for additional cluster characterization including separate beta weights from iE and iC trials (as opposed to the difference score).

#### Drug Effects

To identify regions showing activity alterations linked with drug administration during acute nicotine abstinence, we assessed BOLD signal change following iC feedback, which yielded robust smoker versus nonsmoker differences above, in a GROUP * PATCH * PILL linear mixed-effects framework. Although we did not observe any regions showing GROUP * PATCH * PILL or PATCH * PILL interactions, we detected regions demonstrating significant GROUP * PATCH (ventromedial prefrontal cortex [vmPFC] and caudate) and PATCH main effects (cingulate gyrus and a large cluster encompassing the thalamus, habenula, caudate, lentiform nucleus, and insula; **Supplemental Table S4**). A follow-up within-group repeated-measures ANOVA among abstinent smokers indicated that nicotine (versus placebo) administration was associated with reduced activation in the habenula, bilateral caudate, and cingulate gyrus, (nicotine < placebo) as well as reduced deactivation in the vmPFC following positive feedback (nicotine > placebo) (Fig. 5A, **Supplemental Table S5**). We extracted *β* coefficients from these ROIs for both positive and negative feedback trials across all sessions and participants for qualitative graphical (to avoid a circular analysis [60]) or quantitative statistical examination. Qualitative assessment of activity linked with positive (iC) feedback indicated that nicotine-induced alterations in the habenula were observed among smokers, but not nonsmokers (Fig. 5B). Quantitative statistical assessment of activity linked with negative (iE) feedback lead to the same interpretation (Fig. 5C). This was indicated by a significant GROUP * PATCH interaction (*F*[1,39] = 5.7, *p* = 0.02), which was followed by within-groups repeated-measures ANOVAs identifying a nicotine-induced reduction of habenula activity among smokers (PATCH main effect: *F*[1,21] = 24.6, *p* < 0.001), but not nonsmokers (PATCH main effect: *F*[1,18] = 1.5, *p* = 0.2). Allowing for comprehensive interpretation of pharmacological effects, activity from these ROIs following both positive and negative feedback across all 6 sessions for both smokers and nonsmokers can be found in **Supplemental Fig. S5**. Ancillary ROI-based analyses utilizing anatomically-defined left and right habenula locations further supported the interpretation that nicotine reduced habenula activity following positive and negative feedback among smokers, but not nonsmokers (**Supplemental Fig. S6**).

**Figure 5.**
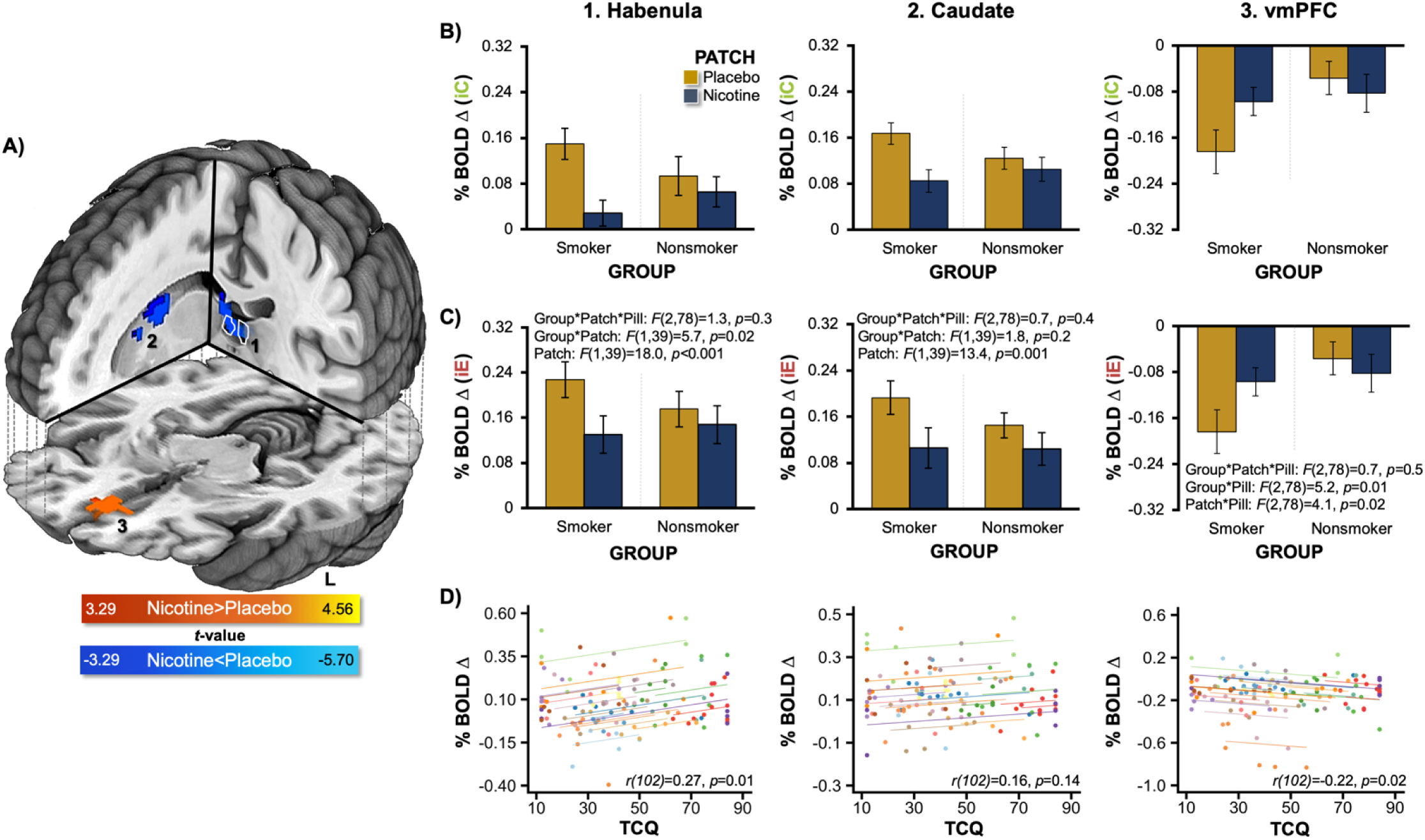
Drug effect: Nicotine-induced alterations following both positive (iC) and negative (iE) feedback. <u>(**A**)</u> Nicotine administration to smokers was associated with reduced activation in the habenula (outlined in white), left and right caudate, and cingulate gyrus as well as reduced deactivation of the vmPFC. These regions were identified when considering the PATCH main effect in a within-group PATCH * PILL repeated-measures analysis among smokers. To indicate regional direction of change, a general linear *t*-test (GLT) contrast was included in the model (cold colors = nicotine < placebo, warm colors = nicotine > placebo, *p_corrected_* < 0.05). <u>(**B**)</u> Qualitative inspection of mean BOLD signal change from select regions indicated that nicotine-induced alterations following positive (iC) feedback were observed among smokers, but not nonsmokers in the 1. habenula 2. right caudate, and 3. vmPFC. <u>(**C**)</u> Selective statistical assessment of mean BOLD signal change from select regions indicated nicotine-induced alterations following negative (iE) feedback were observed among smokers, but not nonsmokers particularly in the habenula. This was indicated by a significant GROUP * PATCH interaction (*p* = 0.02) identified within a GROUP * PATCH * PILL mixed-effects ANOVA. **<u>(D)</u>** Repeated-measures correlations (RMcorr) assessing the within-individual relationships between brain activity and tobacco craving questionnaire (TCQ) total scores, indicated that smokers’ habenula (*p*_Bonferonni-corrected_ = 0.01) and vmPFC (*p*_Bonferronni-corrected_ = 0.02) activity following positive feedback were linked with craving. Indicative of regional specificity, a similar within-individual relationship was not observed when considering caudate activity (*p* = 0.14). In these RMcorr scatter plots, colored circles designate observations from the same smoker (at different sessions) and colored parallel lines designate the RMcorr fits common across each participant. See **Supplemental Table S5** for cluster coordinates.

Consistent with a link to a state-level measure of withdrawal status, a repeated-measures correlation (RMcorr) assessment indicated that higher session-specific habenula activity within an individual smoker was associated with higher state-levels of tobacco craving assessed during that same session (*r*[102] = 0.27, *p* = 0.01, Fig. 5D). Additional exploratory analyses across all participants suggested that higher session-specific habenula activity within an individual participant also correlated with higher state-level social anhedonia assessed that same session (*r*[193] = 0.16, *p* = 0.03, **Supplemental Fig. S7**).

## DISCUSSION

While the ventral striatum, ACC, and insula are regarded as key constituents in the neurocircuitry of addiction [8, 9], emerging preclinical evidence also implicates the habenula as a contributor to negative reinforcement mechanisms perpetuating nicotine use, particularly during the early stages of cessation [33–36]. In the context of a pharmacological fMRI study, we utilized a performance feedback task previously shown to differentially activate these brain regions [45] and delineated functional alterations both as a function of a *chronic* smoking history (smokers vs. nonsmokers) and as a function of drug administration (nicotine, varenicline, vs. placebo) following *acute* smoking abstinence (∼14 hours). Regarding task effects, we observed increased activity following negative feedback in the habenula, ACC, and bilateral anterior insula, as well as increased activity following positive feedback in the bilateral ventral striatum. Regarding group effects, smokers (vs. nonsmokers) showed reduced ventral striatal responsivity to positive feedback, an alteration that was not alleviated by drug administration, but rather was associated with higher *trait-levels* of addiction severity among smokers and elevated self-reported negative affect across all participants. Regarding drug effects, nicotine (vs. placebo) decreased habenula activity following positive and negative feedback in overnight abstinent smokers (but not nonsmokers) with greater habenula activity associated with elevated *state-levels* of tobacco craving among smokers and elevated social anhedonia across all participants. Taken together, these results highlight a dissociation between functional brain alterations underlying two facets of tobacco use disorder, namely, those linked with trait dependence severity (addiction-related) and those linked with state pharmacologic factors (withdrawal-related).

### Task Performance Measures Served as a Drug-Manipulation Check

Behavioral results demonstrated that nicotine and varenicline administration augmented gross task-based attention among smokers and nonsmokers. Specifically, both drugs reduced the number of momentary attentional lapses (i.e., *no response* trials) as indicated by significant within-group PATCH * PILL interactions. This interaction pattern is generally consistent with the known pharmacodynamic actions of nicotine (full agonist) and varenicline (partial agonist) at ⍰4β2 nAChRs [61] and similar to previous observations when assessing heart rate [62], other behavioral measures [22, 62, 63], as well as task-based [15, 22, 62] and resting-state fMRI measures [64], all within the current experimental design and cohort. Performance deficits among cigarette smokers [65] and associated nicotine-induced enhancement manifest in multiple domains, particularly when considering aspects of attention [29]. Whereas nicotine-induced performance enhancement among short-term abstinent smokers represents acute withdrawal reversal, such effects among nonsmokers likely reflects attentional enhancement independent of withdrawal relief [66]. Performance enhancement was less pronounced among our sample of nonsmokers likely due to floor/ceiling effects and the minimal dynamic range for drug administration to further augment performance. These behavioral alterations following nicotine and varenicline administration served as a pharmacologic-manipulation check confirming that these drugs yielded an overt response in both participant cohorts.

### Replication of Neuroimaging Task Effects

When assessing task-related brain activity across all participants, we largely replicated the original implementation of the task that utilized healthy individuals [45]. Specifically, we observed increased habenula, ACC, and bilateral insula activity following negative feedback and increased bilateral nucleus accumbens activity following positive feedback. Given renewed interest in the reproducibility of psychological and, in particular, neuroimaging results [67–69], replication of these task-based outcomes within a larger sample than that from the original report [45] and across multiple scanning sessions is noteworthy. In addition, our results extend the body of extant literature on human habenula function [34, 55, 56, 70] by providing further evidence for the region’s role in reward processing. Specifically, we explored the interrelation of brain activity between pairs of task-related regions (i.e., correlations between feedback-related activity from the habenula, insula, ACC, and striatum). We observed that as the responsivity of the habenula increased to negative feedback across participants, the insula’s and ACC’s responsivity to negative feedback also increased, while conversely, the ventral striatum’s responsivity to positive feedback decreased. These exploratory correlational outcomes are generally consistent with the habenula’s role in negative outcome processing [45, 55], within a larger negative outcome processing neurocircuitry [54, 70], and in inhibiting activity of DAergic neurons which project to the ventral striatum [46, 58].

### Striatal Function and Trait-Level Addiction Severity (Group Effects)

Overnight abstinent cigarette smokers, relative to nonsmokers, had lower neural responses to positive feedback in the bilateral ventral striatum. This finding is consistent with accumulating neuroimaging evidence suggesting that dysregulated reward processing among chronic smokers is mediated by neuroadaptations (and/or preexisting vulnerabilities) within the striatum that manifest as both hypersensitivity to drug-related reward (e.g., cues) [23] and hyposensitivity to monetary rewards [15, 17, 22, 26, 32]. Another noteworthy aspect of this study is that the current task utilized non-monetary feedback (i.e., emojis) to inform participants about trial outcomes, thereby speaking to the generalizability of striatal dysfunction beyond the anticipation and receipt of monetary rewards. Critically, smokers’ blunted striatal responses to positive feedback were not altered by nicotine or varenicline, but rather were correlated with addiction severity (FTND) scores. This relationship between blunted striatal activity and increased trait-levels of addiction severity is largely consistent with our previous observations within the same experimental design, but utilizing a probabilistic reversal learning (PRL) task to probe MCL circuitry [22]. In that report [22], we observed hypoactivation within the bilateral dorsal striatum and dorsal ACC (dACC) following the receipt of monetary rewards among abstinent smokers versus nonsmokers, a deficit that was more pronounced with increasing FTND scores. Although speculative, we suggest that the ventral (current report) versus dorsal (previous report [22]) distinction may reflect differences in the cognitive demands of the tasks employed and the dissociation of dorsal striatal circuitry associated with goal-directed *learning* from ventral striatal circuitry associated with goal-directed *performance* [71]. More specifically, the PRL task [22] involved deciding when to change one’s behavior versus when to maintain a previous behavior, thereby probing dorsal striatal functioning, whereas the current performance feedback task involves less of a learning component and thereby serves more as a probe of ventral striatal functioning. Moving beyond activity within circumscribed regions, additional evidence also implicates the functional connectivity between the striatum and dACC as similarly reflecting addiction severity [72, 73]. As such, we have previously proposed that altered activity within the striatum and/or its functional connectivity with the dACC is linked with trait-level addiction severity as such brain measures are correlated with FTND scores and are not impacted by acute nAChR agonist administration [32]. The current results are consistent with this position and suggest that classes of pharmacologic agents other than nAChR agonists may be needed to target this facet of tobacco use disorder.

### Habenular Function and State-Level Pharmacologic Factors (Drug Effects)

In contrast to ventral striatal activity, habenular activity following both positive and negative feedback *was* modulated by acute nicotine (vs. placebo) administration among abstinent smokers. This finding is consistent with recent theorizing regarding the habenula’s critical role in the development of addiction and mediating the transition from positive to negative reinforcement mechanisms perpetuating drug use [34]. Whereas positive reinforcement relates to learning via the receipt of outcomes engendering a positive state (rewards), negative reinforcement relates to learning via the alleviation of outcomes engendering a negative state (removal of aversive stimuli). In the context of abstinent cigarette smokers, such negative states may include aversive nicotine withdrawal symptoms, including tobacco craving, anhedonia, irritability, and deficits in brain systems linked with reward processing. Indeed, emerging preclinical evidence suggests that the habenula contributes to negative reinforcement mechanisms perpetuating nicotine use including withdrawal-induced anhedonia, anxiety, and depression [33–36].

Noteworthy, we provide novel empirical evidence that elevated habenula activity during acute smoking abstinence can be ameliorated by NRT. Nicotine appeared to down-regulate habenula activity in a “trial non-specific” fashion (i.e., following both positive and negative outcomes) among abstinent smokers during performance feedback. From a different perspective, acute nicotine abstinence was associated with up-regulated habenula activity. Whereas DA neurons increase firing to stimuli predicting positive outcomes and are inhibited by stimuli predicting negative outcomes, habenula neurons show the opposite pattern and increase firing to negative-outcome-predicting stimuli and decrease firing to positive-outcome-predicting stimuli [46]. We speculate that up-regulated habenula activity during acute abstinence may contribute to at least two commonly reported withdrawal symptoms, namely irritability and anhedonia. Operationalizing irritability as enhanced responsivity to negative outcomes, increased habenula activity following negative feedback may be a neurobiological manifestation of this withdrawal symptom. Such a perspective is consistent with the habenula being a critical component of error-related neurocircuitry [56]. Operationalizing anhedonia as attenuated responsivity to a positive outcome, increased habenula activity following positive feedback may, in part, contribute to a hypodopaminergic state and anhedonia. Indeed, when considering all participants, we observed a positive correlation between greater habenula activity following positive feedback and higher state-levels of self-reported social anhedonia. Unfortunately, we did not utilize a validated, multi-item self-report assessment of irritability to provide empirical support for a potential link between elevated habenula activity following negative feedback and higher levels of irritability

Linking such brain alterations with a clinically-relevant construct, we observed that the higher an individual smoker’s session-specific habenula activity was, the greater their state-level of tobacco craving was during that same scanning visit. Habenular structural and functional alterations have previously been linked with major depressive disorder [47, 74–76], a condition in which smokers are overrepresented [77] and in which social anhedonia has been regarded as an endophenotype [78, 79]. Consistent with these prior observations, we observed that the higher an individual participant’s session-specific habenula activity was, the greater their self-reported social anhedonia also was during that same scanning visit. As such, we propose that habenular activity is linked with the state of nicotine withdrawal, given that these brain measures were modulated by NRT in abstinent smokers, unaltered by nicotine in nonsmokers, and linked with self-reported tobacco craving and social anhedonia.

### Limitations

While our experimental design involving drug and placebo administration to smokers and nonsmokers allowed us to characterize functional alterations as both a function of chronic smoking and acute nicotine withdrawal, methodical limitations should be considered. First, the small size of the habenula (∼30mm^3^, [52]) generally limits its assessment in human fMRI studies utilizing “standard” EPI protocols (e.g., re-sampled 3mm isotropic voxels). Given individual variation in habenula locations even after normalization to standard space, smoothing of functional data was a necessary step for voxel-wise group-level analyses [52]. Although we applied a minimal amount of spatial smoothing (smoothed to 7mm FWHM), given the habenula’s small size we cannot rule out the possibility that signals within the anatomical habenula were potentially contaminated by signals from surrounding regions/structures. That said, the regions immediately adjacent to the habenula that, due to partial volume effects, could have contributed to the signal measured (e.g. mediodorsal thalamus) are not known to have functional properties consistent with that identified for the habenula in preclinical models [57, 80]. Related, given the habenula’s adjacency to the ventricles, BOLD signal in this location may be susceptible to physiological artifacts (e.g., pulsation, respiration) which are a source of noise in fMRI data [81]. While we did not collect peripheral recordings of heart rate or respiration to minimize the already high participant burden of this study, we acknowledge that the inclusion of such measures as regressors of no interest could potentially improve assessments of this region. Continued improvements in equipment hardware and continued development of open-source toolboxes [82] will increase the more widespread adoption of physiological noise reduction. These limitations may be at least partially circumvented utilizing higher resolution imaging techniques achieved at 7T [83] or at 3T [44, 47] yielding voxel resolution in the 1mm-2mm isotropic range as well as enhanced preprocessing for physiological noise reduction. Second, we acknowledge the functional specialization of the medial and lateral subdivisions of the habenula identified in preclinical work such that medial aspects possessing higher densities of nAChRs have been primarily linked with aversive responses to high nicotine doses [38, 39] and the lateral habenula having been primarily linked with reward processing, aversive stimuli, and emotions [70]. However, given the resolution of our fMRI data we are not able to speak to such sub-regional implications. Third, to ensure sufficient power needed to detect higher-level group and drug effects on task-based brain activity, we employed a small volume corrected mask based on *a priori* hypotheses of brain regions implicated in nicotine withdrawal. However, group and drug effects in brain regions extending outside of our *a priori* mask may provide additional insight into neurobiological processes relevant to smoking-related behaviors. Finally, due to sample size limitations, the effect of biological sex on these outcomes were not directly assessed despite emerging evidence of differential brain activity linked to nicotine withdrawal in females and males [84].

### Conclusions

This study highlights a dissociation between neurobiological processes linked with trait-level addiction severity and those linked with state-level nicotine withdrawal. Utilizing a performance feedback task, we replicated differential patterns of brain activity in the habenula, insula, ACC, and ventral striatum following positive and negative feedback. Abstinent smokers showed reduced striatal responsivity to positive feedback, an alteration that was not ameliorated by administration of two commonly used pharmacologic smoking cessation aids, but rather was correlated with severity of nicotine addiction. Conversely, nicotine administration decreased habenula activity following positive feedback among abstinent smokers and such activity was correlated with self-reported tobacco craving and social anhedonia. These outcomes provide novel evidence linking human habenula function with state-like aspects of tobacco use disorder. Delineating the brain mechanisms perpetuating cigarette smoking that are, and are not, mitigated by NRT may expedite development of improved smoking cessation interventions. Specifically, these outcomes provide insight into why common pharmacologic interventions fail for most smokers. That is, nicotinic drug administration did not appear to impact trait-levelbs dysregulated striatal activity following positive feedback. Interventions simultaneously targeting both trait-like addiction-related and state-like withdrawal-related facets of tobacco use disorder may improve cessation outcomes.

## MATERIALS AND METHODS

### Participants

A total of 24 non-treatment seeking daily cigarette smokers (≥10 cigarettes per day for >2 years, 12 females) and 20 nonsmokers (no smoking within the last 2 years and no lifetime history of daily smoking, 10 females) completed the study. All 44 participants were right-handed, 18-55 years of age, and were recruited at the National Institute on Drug Abuse-Intramural Research Program (NIDA-IRP) in Baltimore, MD. Participants were healthy with no reported history of drug dependence (other than nicotine in smokers), neurological or psychiatric disorders, cardiovascular or renal impairments, diabetes, or MRI contraindications. Cigarette smokers were 36 ± 10 years old (mean ± SD), reported daily smoking for 18 ± 11 years, smoked 18 ± 8 cigarettes per day, and were moderately nicotine dependent (FTND scores: 5 ± 2, **Supplemental Table S1**). As nonsmokers were younger (30 ± 7 years old) than smokers, we considered the influence of age when comparing groups. The results and interpretations when including age as a covariate in between-group behavioral and neuroimaging assessments were the same as when age was not included. All participants’ data were utilized in behavioral assessments, however for imaging assessments, fMRI data from 2 male smokers and 1 male nonsmoker were excluded due to excessive head motion during scanning (criteria described below). We obtained written informed consent in accordance with the NIDA-IRP Institutional Review Board and volunteers were compensated for participation. This study was registered at clinicaltrials.gov (NCT00830739).

### Experimental Design and Procedures

We collected self-report questionnaire, behavioral task performance, and MRI data from participants in the context of a within-subject, double-blind, placebo-controlled, crossover study, involving two drugs: transdermal nicotine (NicoDerm CQ, GlaxoSmithKline) and oral varenicline (Chantix, Pfizer). The 6-8 week study duration involved a total of 9 visits, each on different days (1 orientation, 2 neurocognitive, and 6 neuroimaging visits; Fig. 1A) [15, 22, 62, 64, 85]. During the orientation visit, participants gave informed consent, completed baseline self-report measures, and received training on multiple neuroimaging tasks (one described herein). Task instructions were initially explained at a bench computer and then practice was completed in a mock MRI scanner. During the neurocognitive visits, participants completed self-report measures and behavioral testing (not described further). At three time points during a varenicline administration regimen (PILL factor: pre-pill [baseline] vs. placebo vs. varenicline), participants completed MRI scanning on two occasions, once while wearing a nicotine patch and once while wearing a placebo patch (PATCH factor). Following two initial pre-pill imaging sessions, each participant underwent ∼17 days of varenicline and placebo pill administration and completed nicotine and placebo patch visits towards the end of both pill periods. During each of the 6 neuroimaging days, participants completed two MRI sessions lasting ∼2 hours each (an AM and a PM session). The AM session began ∼2-2.5 h after patch application. During this session, participants completed a Flanker task [63], a performance feedback task (described below), and a response inhibition task. The performance feedback task was completed ∼45 min after scanner entry and 2.5-3.5 h after patch application.

Varenicline was administered according to standard guidelines beginning with a once-daily dose of 0.5 mg on days 1-3, 0.5 mg twice daily on days 4-7, and increased to 1 mg twice daily on days 8-17. (https://www.pfizer.com/products/product-detail/chantix). Active varenicline and placebo pills were packaged to be identical in appearance. Scanning sessions occurred at the end of each regimen (varenicline 17.0 ± 4.2 days; placebo pill 16.5 ± 3.4 days). Pill administration periods were not separated by a washout interval. For participants with a placebo regimen that followed the varenicline regimen, carryover effects were assumed negligible given the ∼24-hour elimination half-life of varenicline and the fact that placebo-pill scanning sessions and active varenicline-pill scanning sessions were separated by more than 2 weeks. Adherence to the pill regimen was confirmed by pill count and participant self-report the morning of each MRI visit; data on medication adherence have been reported elsewhere [63]. Pill regimen side effects and adherence were monitored by regular telephone assessments and at in-person visits.

For each of the pill conditions, nicotine or placebo patches were applied to the upper back at the start of fMRI visits (separated by 2.9 ± 1.7 days). The patch was worn for the duration of each 9-hour neuroimaging visit. Pharmacokinetic data indicate plasma nicotine concentrations reach a peak within 2-4 h after patch application, remain relatively stable for the next 4-6 h, and then gradually decrease beginning ∼8-10 h post-patch [86]. As such, data collection occurred within a 2-9 h post-patch window associated with steady plasma nicotine levels. All nonsmokers were administered 7 mg nicotine patches. For smokers, a multiple dosing strategy was employed to match daily nicotine intake: 21 mg (10-15 cigs/day; n = 11), 28 mg (16-20 cigs/day; n = 9), 35 mg (21-25 cigs/day; n = 1), and 42 mg (> 25 cigs/day; n = 3).

Overnight abstinence was required on neuroimaging days. Smokers were instructed to have their last cigarette 12 hours before their scheduled arrival. All participants were instructed to abstain from alcohol for 24 hours, and to moderate caffeine intake for 12 hours. Upon arrival, all participants were tested for recent drug and alcohol use, and for expired carbon monoxide (CO) levels. A CO guideline of less than 15 parts per million (ppm) was used to verify smoking abstinence. Indicative of compliance, smokers’ CO levels were lower at neuroimaging visits (7.1 ± 2.6 ppm) in comparison to the orientation and neurocognitive visits which did not require abstinence (18.9 ± 8.9 ppm; *t*[23] = −8.2, *p* < 0.001). Nonsmokers’ CO levels did not differ between such visits (abstinence required: 1.9 ± 0.3 ppm; not required: 1.8 ± 0.4 ppm; *p* = 0.2).

### Self-report measures

We assessed clinically-relevant constructs (addiction severity, tobacco craving, affect, and anhedonia) using previously validated and commonly employed self-report instruments. In the context of cigarette smoking, addiction severity is typically quantified with the 6-item FTND [87]. FTND scores are highly heritable [88] and routinely utilized as a primary phenotype in studies linking smoking behaviors with nicotinic acetylcholine receptor and other genetic variants [89]. As such, we conceptualized FTND scores (range: 1-10) collected during the orientation visit as a *trait-level* measure of nicotine addiction severity. Additionally, smoker’s tobacco cravings were assessed each neuroimaging visit with the 12-item, short-form of the Tobacco Craving Questionnaire (TCQ) [90]. TCQ items are rated on a 7-point scale (strongly disagree to strongly agree) and yield a total score (range: 12-84) and four subscale scores (emotionality, expectancy, compulsivity, and purposefulness). We focused on the TCQ total score which we conceptualized as a *state-level* measure of nicotine withdrawal status. All participants also completed instruments to assess affect and anhedonia at each neuroimaging visit. Specifically, participants completed the Positive and Negative Affect Schedule (PANAS) [91] which is composed of two 10-item scales assessing positive affect (active, alert, attentive, determined, enthusiastic, excited, inspired, interested, proud, strong) and negative affect (afraid, ashamed, distressed, guilty, hostile, irritable, jittery, nervous, scared, upset). PANAS items were rated on a 5-point scale (not at all/very little to extremely) yielding positive and negative affect scores with a possible range from 10 to 50. Participants also completed the Revised Social Anhedonia Scale (RSAS) [92, 93] which is a 40-item true-false questionnaire that measures pleasure (or lack thereof) derived from interpersonal sources [94]. RSAS items are dichotomously scored and total scores range from 0 to 40 such that higher values indicate greater levels of anhedonia. Elevated social anhedonia is common among individuals diagnosed with depression and schizophrenia [94], two neuropsychiatric conditions linked with aberrant habenula function [57, 95–98] and in which smokers are over-represented [99–101]. The TCQ (smokers only), PANAS, and RSAS were collected each neuroimaging visit ∼1.5 hours after completion of the performance feedback task.

### Performance feedback task

At each neuroimaging visit, participants performed a positive and negative performance feedback task, called the motion prediction task, previously shown to differentially activate the habenula, ACC, insula, and ventral striatum [45]. The participants’ objective was to predict which of two moving balls, starting from different locations and traveling at different speeds, would be the first to reach a finish line after viewing a brief clip of the balls’ motion (Fig. 1B). Each trial began with a variable fixation interval (3500-4500ms) which was followed by the appearance of the finish line (500ms) to signify an upcoming motion event. Participants then viewed a short sequence (1400ms) of the two balls traveling at different speeds from different starting locations on the left side of the screen towards the finish line on the right side of the screen. Still far from the finish line, the balls disappeared and the question “Which Ball?” was presented (1350ms). Participants indicated their prediction/decision via a right-handed button press with either the index (ball 1) or middle finger (ball 2) during this response window. After a fixed delay (750ms), performance feedback (emojis) about the correctness of the participants’ prediction was presented (1000ms).

Performance feedback was delivered to participants in a two-factor, FEEDBACK (informative [i] vs. non-informative [n]) * RESPONSE (correct [C] vs. error [E]), fashion. Under informative feedback, participants always received positive feedback (i.e., a happy emoji) following a correct response (iC) and always received negative feedback (i.e., a sad emoji) following an erroneous response (iE). Under noninformative feedback, participants received no information on whether they gave a correct or erroneous response as the same feedback was presented on both nC and nE trials (i.e., an ambiguous emoji). Informative feedback was presented on 73% of trials and noninformative feedback on 27%. This feedback schedule allowed us to determine which task-related aspect more robustly influenced brain activity, namely, the type of feedback (positive vs. negative) or the type of response (correct vs. error). Task-related blood oxygenation level-dependent (BOLD) signal change was anticipated to be robustly modulated on error versus correct trials when followed by informative feedback. If participants failed to respond during the designated response window, they received a fifth type of feedback (i.e., a confused emoji) for these *no response* trials.

Task difficulty, operationalized as the time difference between the two balls’ arrival at the finish line, was dynamically manipulated based on each participant’s behavior to maintain error rates at ∼35% (i.e., an error rate between 0.3 and 0.4 over a sliding window). Specifically, over a 10-trial sliding window, if error rates were below 0.3, task difficulty was increased and if the rate was above 0.4, task difficulty was decreased. This task feature was intended to induce participant uncertainty about trial performance until feedback delivery (i.e., to mitigate the self-detection of errors). While the dynamic difficulty manipulation was intended to consistently maintain error rates across participants and across the 6 neuroimaging visits, one behavioral aspect not under the control of the task’s adaptive algorithm was the number of *no response* trials (errors of omission), which reflect momentary lapses of attention. For each *no response* trial encountered, the adaptive algorithm reduced the targeted number of *correct* trials by one. As such, the percent of *no response* and *correct* trials were inversely related, demonstrated more variability across participants and sessions than that for *error* trials, and thus provided a convenient behavioral metric to assess group- and drug-related effects on these two dependent variables which we interpreted as gross behavioral measures of attention. Given that the percent of *no response* and *correct* trials were ‘mirror images’ of each other, only *no response* trial results are presented. Task presentation and behavioral performance recording were controlled by E-prime software (v1.2, Psychology Software Tools). Participants completed a total of 240 trials in four 9-minute runs with short rest periods between runs 1-2 and 3-4 and a longer break between runs 2-3 (during which a T1-weighted structural MRI was collected and participants were instructed to relax while staying as still as possible).

### Behavioral measures: Statistical analysis

The primary behavioral performance measures considered from the task were response times (RT) and the percent of *no response* trials. To replicate outcomes from the original task implementation [7] and to assess task engagement, we analyzed RT data in a FEEDBACK (informative [i] vs. non-informative [n]) * RESPONSE (correct [C] vs. error [E]) repeated-measures ANOVA using SPSS (v23, Chicago, IL). As this analysis focused on the *overall task effect* (as opposed to session-specific fluctuations), each participant’s RT data were first averaged across all 6 neuroimaging visits (i.e., collapsed across session) separately for each trial type. Main and interaction effects were considered. To characterize *group effects* and *drug effects* on an objective measure of attention, we analyzed the percent of *no response* trials in a mixed-effects ANOVA including factors for GROUP (between-subjects: smokers vs. nonsmokers), PATCH (within-subjects: nicotine vs. placebo), and PILL (within-subjects: pre-pill, varenicline, vs. placebo). Main and interaction effects were considered and significant interactions involving the GROUP factor and/or a PATCH main effect were followed by within-group, repeated-measures ANOVAs including PATCH and PILL as factors. Geisser-Greenhouse corrections for violations of sphericity were utilized when considering effects involving the PILL factor (3 levels). Pairwise follow-up *t*-tests comparing nicotine versus placebo PATCH differences (in the absence of varenicline under placebo pill conditions) and assessing varenicline versus placebo PILL differences (in the absence of nicotine under placebo patch conditions) were Bonferroni-corrected. All participants (*N* = 44) sufficiently performed the task and contributed behavioral data to these analyses.

### MRI data acquisition and analysis

MRI data were collected on a Siemens 3T Magnetom Allegra scanner (Erlangen, Germany). During task performance, 33 slices (5mm thick) were obtained in the sagittal plane using a T2*-weighted, single-shot, gradient-echo, echo-planar imaging (EPI) sequence sensitive to BOLD effects (272 volumes/run, repetition time = 2,000ms [332ms delay], echo time = 27ms, flip angle = 80°, field of view = 220 mm in a 64 x 64 matrix). As simultaneous EEG data were also acquired (not discussed further), sagittal images were collected with a delay between volume acquisitions to aid scanner-artifact removal from EEG recordings. T1-weighted structural images were obtained using a magnetization prepared rapid gradient-echo sequence (MPRAGE: repetition time = 2,500ms; echo time = 4.38ms; flip angle = 8°; voxel size = 1mm^3^).

Neuroimaging data were preprocessed and analyzed using AFNI (version 16.3.18; http://afni.nimh.nih.gov/afni/). Functional images were slice-time and motion corrected, registered to the anatomical volume, normalized into Talairach space (3mm isotropic voxels, 27 *µL*), and spatially blurred to 7mm full-width at half-maximum (AFNI’s 3dBlurToFWHM). Following motion correction, functional volumes/frames with > 0.35mm Euclidean norm mean displacement were censored along with the immediately adjacent volumes. Those volumes with fewer than three contiguous uncensored neighbors were also flagged for censoring. Individual scanning sessions were excluded from imaging analyses if more than 25% of the functional volumes were motion censored. This resulted in the exclusion of 3 male subjects (2 smokers, 1 nonsmoker) from further analyses given that a majority of their 6 neuroimaging sessions were flagged as high motion. Six additional subjects had between 1 to 3 scanning sessions discarded from further analysis due to motion. As such, 41 participants (22 smokers, 19 nonsmokers) contributed neuroimaging data to these analyses.

Functional time series were normalized to percent signal change and submitted to voxel-wise multiple regression. In the first-level general linear model (GLM), we included five separate task-related regressors as impulse functions time-locked to the onset of feedback presentation (iC, iE, nC, nE, and *no response* trials). These regressors were convolved with a model hemodynamic response (gamma) function and its temporal derivatives. Regressors of no interest also included in the GLM were the six motion-correction parameters as well as fourth-order polynomial regressors to capture residual head motion and baseline trends in the BOLD signal, respectively. As the time series were scaled by the voxel-wise mean, resulting regression (*β*) coefficients, calculated per regressor, participant, and session, were interpreted as an approximation of percent BOLD signal change (% BOLD Δ) [102] from the implicit baseline.

#### Task Effects

To replicate outcomes from the original task implementation [45] and to identify regions showing differential activity following performance feedback, we calculated contrast images between negative and positive feedback (iE – iC betas). Individual participant’s contrast images were first averaged across all sessions and then submitted to a group-level, whole-brain, one-sample *t*-test (2-tailed, 3dTtest++). The resulting statistical maps were thresholded correcting for family-wise error at *p_corrected_* < 0.001 (*p_voxel-wise_* < 0.0001, cluster-extent: 15 voxels, 51,704 voxels in the whole-brain mask, 3dClustSim with the spatial autocorrelation function correction [103]). For graphical examination and follow-up analyses, we extracted the mean *β* coefficients associated with specific task events (iE, iC, nE, nC) and the [iE – iC] contrast values from the identified task-related clusters/regions of interest (ROIs) by averaging across all voxels within each separate ROI. We performed exploratory bivariate Pearson’s correlations between the contrast values (iE – iC betas) from pairs of identified ROIs to examine the relationship between the feedback-related responsivity of the habenula, insula, and ventral striatum. These exploratory correlation analyses were not Bonferronni-corrected for multiple comparisons.

#### Group effects

To characterize functional alterations linked with a *chronic* smoking history, we compared session-averaged iE – iC contrast images between smokers and nonsmokers in a 2-tailed independent samples *t*-test (3dTtest++). Given our *a priori* focus on brain regions previously linked with addiction, nicotine withdrawal, and feedback processing in the mesocorticolimbic circuitry [15, 22, 45, 104], we applied a family-wise error correction (*p_corrected_* < 0.05) within a composite mask of interest consisting of the bilateral nucleus accumbens, caudate, putamen, pallidum, anterior insula, anterior cingulate, middle anterior cingulate, subcallosal cingulate, and habenula regions (**Supplemental Fig. S3**). All anatomical masks came from the Desai maximum probability map [105], with the exception of the habenula mask which was defined by placing 3mm radius spheres around left and right hemisphere coordinates previously reported for the motion prediction task [45]. Within the composite mask, statistical maps were thresholded correcting for family-wise error at *p_corrected_* < 0.05 (*p_voxel-wise_* < 0.001, cluster-extent: 9 voxels, 7,684 voxels in composite mask). For graphical examination and follow-up analyses, we extracted the mean contrast values (iE – iC betas) as well as the individual *β* coefficients for iE and iC trials from those ROIs showing smoker versus nonsmoker differences. To link altered brain activity among smokers with a clinically-relevant construct, we conducted *a priori* hypothesized Pearson’s correlations between the [iE – iC] contrast values from identified ROIs and smokers’ FTND scores which reflect trait-levels of addiction severity. The *p*-values resulting from these *a priori* correlation analyses were Bonferronni-corrected (*n* = 3; the number of ROIs considered). To further link feedback-related brain activity from all participants with self-reported affect, we conducted *post hoc* exploratory correlations between [iE – iC] contrast values and each participants’ PANAS scores. Given we were interested in trait-levels of affect (paralleling the session-averaged, trait-level assessment of brain activity), we computed a session-averaged metric of positive and negative affect. Specifically, participants completed the PANAS at each study visit and we were interested in an individual’s general level of affect as opposed to session-to-session fluctuations. These exploratory correlation analyses were not Bonferronni-corrected for multiple comparisons.

#### Drug effects

To delineate functional alterations linked with *acute* drug administration, we assessed *β* coefficient images from iC (positive) feedback trials in a linear mixed-effects (LME) framework (3dLME, v1.9.8) [106] including factors for GROUP (between-subjects: smoker vs. nonsmoker), PATCH (within-subjects: nicotine vs. placebo), and PILL (within-subjects: pre-pill, varenicline, vs. placebo). The LME framework was used given missing session data associated with our motion censoring criteria described above. We focused on iC feedback trials given that a larger effect size was observed for these trials (relative to iE feedback) when assessing smoker versus nonsmoker differences in the group effects analysis above. Significant interaction effects involving the GROUP factor and/or PATCH main effects were followed by within-group (i.e., smokers) LME analyses including PATCH and PILL as factors. Main and interaction effects were of interest. Within this analysis model, a general linear *t*-test (GLT) was performed comparing nicotine versus placebo patch to visually represent the direction of drug-related alterations (i.e., nicotine < placebo, nicotine > placebo) associated with the PATCH main effect. To maximize the potential for producing generalizable results and minimizing Type I error rates, this LME model utilized a “maximal random effects structure” [107, 108] (i.e., included random intercepts for subjects and random slopes for the PATCH, PILL, and PATCH * PILL effects; -ranEff ‘∼1 + patch + pill + pill*patch’). Statistical maps were thresholded correcting for family-wise error within the composite mask as described above for the group effects analysis. We extracted the *β* coefficients from both the iC and iE trials from those ROIs showing drug-induced effects separately for all 6 drug conditions and across both groups (smokers, nonsmokers) for qualitative graphical (to avoid a circular analysis [60]) and/or selective quantitative statistical examination thereby facilitating comprehensive interpretation of pharmacological effects. While a selective analysis of smokers’ iC *β* coefficients would constituent a circular analysis [60], we extracted these parameter estimates for graphical representation. Given the iE *β* coefficients were unrelated to the statistical used to identify voxels of interest, we assessed parameter estimates from select regions in a GROUP * PATCH * PILL mixed-effects ANOVA using SPSS. Significant interactions involving the GROUP factor and/or PATCH main effectswere followed by within-group, repeated-measures ANOVAs including PATCH and PILL as factors

To link session-to-session fluctuations in brain activity among smokers with a clinically-relevant construct, we conducted *a priori* hypothesized repeated-measures correlations (RMcorr) between session-specific iC beta values from identified ROIs and smokers’ session-specific TCQ scores which reflect a state-level measure of withdrawal status. RMcorr is a statistical procedure implemented in R (v3.5.0) [109] to characterize the within-participant association between two measures assessed on two or more occasions that was common across all participants [110]. The *p*-values resulting from these *a priori* RMcorr analyses were Bonferronni-corrected (*n* = 3; the number of ROIs considered). To further link session-to-session fluctuations in brain activity from all participants with self-reported anhedonia, we conducted *post hoc* exploratory RMcorr between session-specific iC beta values and each participants’ session-specific RSAS scores. These exploratory RMcorr analyses were not corrected for multiple comparisons.

## ACKNOWLDGEMENTS

### Funding Statement

This work was sponsored by the National Institute on Drug abuse, Intramural Research Program, National Institutes of Health, US Department of Health and Human Services (EAS, TJR, BJS, MTS). Co-authors were in part also supported by National Institute on Drug Abuse grant K01DA037819 (MTS), National Institute on Drug Abuse grant R01DA041353 (ARL, MTS, MCR, RP), and National Institute on Minority Health and Health Disparities grant U54MD01239 (sub-project 5378, MTS, JSF). The content is solely the responsibility of the authors and does not necessarily represent the official views of the National Institutes of Health.

### Competing Interests

The authors declare that they have no competing interests.

### Author Contributions

Dr. Sutherland had full access to all the data in the study and takes responsibility for the integrity of the data and accuracy of data analysis. *Study concept and design:* Sutherland, Salmeron, Ross, Stein. *Acquisition, analysis, or interpretation of data*: All authors. *Drafting of the manuscript:* Flannery, Sutherland. *Critical revision of the manuscript for important intellectual content:* All authors. *Statistical analysis*: Flannery, Sutherland. *Obtained Funding:* Stein, Sutherland. *Administrative, technical, or material support:* Riedel, Laird, Ross, Salmeron, Stein, Sutherland. *Study supervision:* Sutherland, Salmeron, Stein.

### Data

All data needed to evaluate the conclusions in the paper are presented in the paper and/or Supplemental Materials. Additional data or materials are available from authors upon request.

