## Supplemental Material for "Habenular and striatal activity during performance feedback are differentially linked with state-like and trait-like aspects of tobacco use disorder"

### SUPPLEMENTAL CONTENT

#### TABLES

- Table S1: Participant demographic and smoking characteristics (pp. 2)
- Table S2: Task effect cluster coordinates (pp. 3)
- Table S3: Group effect cluster coordinates (pp. 4)
- Table S4: Drug effect cluster coordinates across all participants (pp. 5)
- Table S5: Drug effect cluster coordinates among smokers (pp. 6)

#### FIGURES AND LEGENDS

- Figure S1: Anatomically-defined habenula locations across all participants (pp. 7)
- Figure S2: Task effects: Whole-brain RESPONSE * FEEDBACK interaction analysis (pp. 8).
- Figure S3: *A priori* composite mask of interest (pp. 9)
- Figure S4: Group effects: Additional characterization of regions showing smoker versus nonsmoker differences in feedback responsivity (pp. 10-11)
- Figure S5: Drug effects: Brain activity following both positive (iC) and negative (iE) feedback across all session for smokers and nonsmokers (pp. 12-13).
- Figure S6: Drug-effects: Anatomically-defined left and right habenula ROI analyses assessing nicotine-induced alterations following positive (iC) and negative (iE) feedback (pp. 14-15).
- Figure S7: Drug effects: Additional characterization of regions showing nicotine-induced alterations (pp. 16).

### SUPPLEMENTAL TABLES

**Table S1. Participant demographic and smoking characteristics.**

| | Smokers ($n$ = 24) | Nonsmokers ($n$ = 20) | Group differences |
|---|---|---|---|
| Gender (F/M) | 12 / 12 | 10 / 10 | -- |
| Age | 35.7 ± 9.9 (22-52) | 30.1 ± 7.1 (20-45) | $t(41.3) = -2.1$, $p = 0.04$* |
| IQ | 106.6 ± 11.7 (87-127) | 112.7 ± 11.6 (87-130) | $t(40.6) = 1.7$, $p = 0.1$ |
| Race (AA / C / A / >1) | 6 / 13 / 3 / 2 | 7 / 8 / 3 /2 | $\chi^2(4, 44) = 2.1$, $p = 0.7$ |
| Fagerström score | 5.0 ± 1.9 (2-9) | -- | -- |
| Years daily smoking | 18.0 ± 10.6 (3-39) | -- | -- |
| Cigarettes per day | 17.7 ± 7.9 (10-40) | -- | -- |

Note. Data are expressed as mean ± SD (range). AA: African American, C: Caucasian, A: Asian, >1: more than one race. * $p < 0.05$.

**Table S2. Task effect cluster coordinates.** Regions showing greater activation following negative (iE) or positive (iC) feedback across all participants and conditions.

| | Region | Hemisphere | Center Coordinates (Talairach, LPI) X | Y | Z | Cluster Size (# of Voxels) |
|---|---|---|---|---|---|---|
| **Negative feedback (iE > iC)** | | | | | | |
| 1 | **Anterior insula**/inferior frontal gyrus (IFG) | R | 46 | 14 | 14 | 1137 |
| 2 | **Anterior insula**/inferior frontal gyrus (IFG) | L | -42 | 12 | 12 | 992 |
| 3 | **ACC/Pre-SMA/SMA** (superior, middle & medial frontal gyrus, cigulate gyrus) | R,L | 2 | 18 | 46 | 969 |
| 4 | **Thalamus** (encompassing Habenula) | R,L | 1 | -19 | 0 | 748 |
| 5 | **Middle & superior frontal gyrus** | R | 31 | 45 | 19 | 44 |
| 6 | **Posterior insula** & superior temporal gyrus | L | -36 | -29 | 15 | 33 |
| 7 | **Posterior cingulate** | R,L | 0 | -22 | 30 | 50 |
| 8 | **Superior & middle temporal** gyrus, supramarginal gyrus | R | 49 | -45 | 21 | 642 |
| 9 | **Superior & middle temporal** gyrus | L | -45 | -51 | 30 | 46 |
| 10 | **Middle temporal gyrus** & superior occipital gyrus | R | 38 | -71 | 23 | 19 |
| 11 | **Supramarginal gyrus**, superior temporal gyrus, & inferior parietal lobule | L | -45 | -51 | 30 | 376 |
| 12 | **Fusiform gyrus** | R | 37 | -51 | -16 | 295 |
| 13 | **Fusiform gyrus** | L | 49 | -45 | 21 | 217 |
| 14 | **Precuneus** | R | 10 | -68 | 41 | 37 |
| 15 | **Cerebellum** (uvula, pyramis) | L | -18 | -69 | -25 | 35 |
| **Positive Feedback (iE < iC)** | | | | | | |
| 1 | **Ventral striatum** (nucleus accumbens, caudate) | R | 15 | 10 | -3 | 55 |
| 2 | **Ventral striatum** (nucleus accumbens, caudate) | L | -12 | 10 | -3 | 42 |

NOTE. Voxel size: 3 x 3 x 3 mm (27 μL). X: Left (-), Right (+); Y: Posterior (-), Anterior (+); Z: Inferior (-), Superior (+). Region labels come from the AFNI Talairach daemon atlas. **See main text Fig. 3A** for graphical representation.

**Table S3. Group effect cluster coordinates.** Regions showing smoker versus nonsmoker differences in feedback responsivity ([iE – iC] contrast values).

| | Region | Hemisphere | Center Coordinates (Talairach, LPI) X | Y | Z | Cluster Size (# of Voxels) |
|---|---|---|---|---|---|---|
| **Smokers > Nonsmokers (iE - iC Beta)** | | | | | | |
| 1 | **Ventral striatum** (lentiform nucleus, caudate) | R | 13 | 2 | -1 | 29 |
| 2 | **Ventral striatum** (lentiform nucleus) | L | -12 | 0 | -3 | 21 |
| 3 | **Anterior insula**/inferior frontal gyrus (IFG) | L | -32 | 26 | -3 | 18 |

NOTE. Voxel size: 3 x 3 x 3 mm (27 μL). X: Left (-), Right (+); Y: Posterior (-), Anterior (+); Z: Inferior (-), Superior (+). Region labels come from the AFNI Talairach daemon atlas. See **main text Fig. 4A** for graphical representation.

**Table S4. Drug effect cluster coordinates across all participants.** Regions showing significant effects in a GROUP * PATCH * PILL linear mixed-effects analysis (AFNI's 3dLME) assessing brain activity following positive (iC) feedback.

| | Region | Hemisphere | Center Coordinates (Talairach, LPI) x | y | z | Cluster Size (# of Voxels) |
|---|---|---|---|---|---|---|
| **GROUP x PATCH interaction** | | | | | | |
| 1 | **Ventromedial PFC** (medial frontal gyrus, BA10) | R,L | 0 | 41 | -10 | 21 |
| 2 | **Caudate** (body) | L | -16 | 1 | 16 | 23 |
| **PATCH main effect** | | | | | | |
| 1 | **Thalamus/Caudate/Insula/Lentiform nucleus** | R,L | 4 | -3 | 10 | 506 |
| 2 | **Cingulate gyrus** (medial & superior frontal gyrus) | R,L | 1 | 13 | 42 | 144 |

NOTE. Voxel size: 3 x 3 x 3 mm (27 μL). X: Left (-), Right (+); Y: Posterior (-), Anterior (+); Z: Inferior (-), Superior (+). Region labels come from the AFNI Talairach daemon atlas.

**Table S5. Drug effect cluster coordinates among smokers.** Regions showing a significant main effect of nicotine in a PATCH * PILL linear mixed-effects analysis (AFNI's 3dLME) assessing brain activity following positive (iC) feedback among smokers.

| | Region | Hemisphere | Center Coordinates (Talairach, LPI) | | | Cluster Size |
|---|---|---|---|---|---|---|
| | | | X | Y | Z | (# of Voxels) |
| **PATCH main effect @ Smokers** | | | | | | |
| 1 | **Caudate/Lentiform Nucleus** | L | -14 | -1 | 13 | 98 |
| 2 | **Thalamus** (habenula, medial dorsal nucleus) | R,L | 3 | -25 | 8 | 37 |
| 3 | **Caudate** (body) | R | 15 | 3 | 17 | 36 |
| 4 | **Ventromedial PFC** (medial frontal gyrus, BA10) | R,L | 1 | 40 | -10 | 27 |
| 5 | **Cingulate gyrus** (medial frontal gyrus) | R,L | 4 | 11 | 43 | 15 |

NOTE. Voxel size: 3 x 3 x 3 mm (27 μL). X: Left (-), Right (+); Y: Posterior (-), Anterior (+); Z: Inferior (-), Superior (+). Region labels come from the AFNI Talairach daemon atlas. See **main text Fig. 5A** for graphical representation.

### FIGURES AND LEGENDS

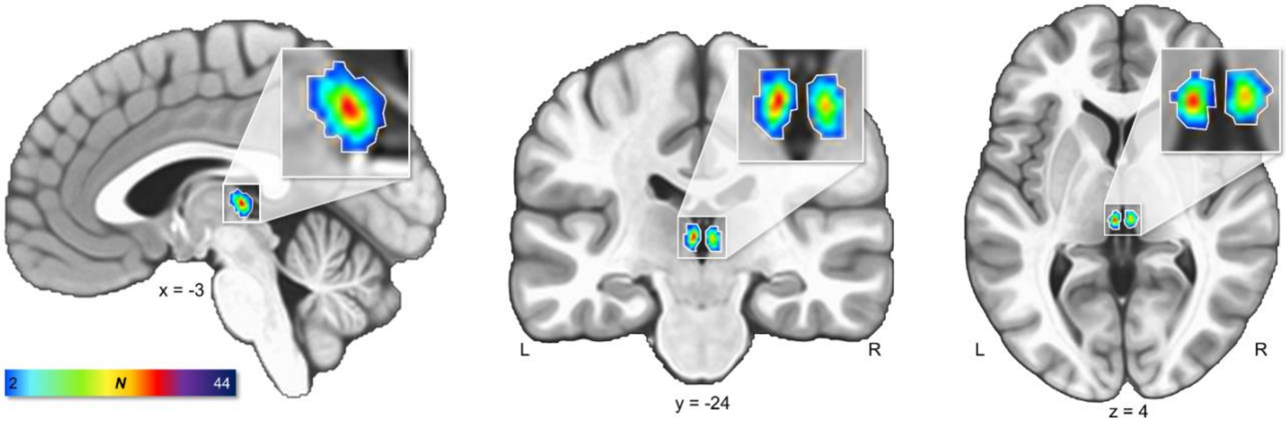


**Figure S1. Anatomically-defined habenula location across all participants.** To provide an anatomical frame of reference for the functional imaging outcomes, we manually defined the habenular complex within each individual participant's T1-weighted structural image. Utilizing anatomical landmarks and visible tissue contrast (detailed below), the left and right habenula were manually traced using AFNI's *Draw Dataset* plugin on all six of each participant's AC-PC aligned and skull-stripped structural images. Habenula-designated voxels were assigned a value of 1 and all other voxels a value of zero within each individual structural volume. These participant- and session-specific habenula masks were summed across all participants (separately for each neuroimaging session) and then registered (normalized) to Talairach space. For each of the 6 sessions, this produced an overlap image with voxel values possibly ranging from 0 to 44, as all participants (24 smokers and 20 nonsmokers) yielded a useable T1-weighted image. Shown above is the overlap map from a single session (i.e., placebo-pill, nicotine-patch session) depicting voxels where a minimum of two participants overlapped. The voxels with the maximum overlap included 29 participants (66% of the sample). The center of mass for the left and right anatomically defined habenular regions were X = -3, Y = -24, Z = 5 and X = 4, Y = -24, Z = 4 in Talairach space, respectively. Given individual variation in post-normalized habenula locations across participants, we created an inclusive anatomical frame of reference by outlining those voxels with at least an overlap of 2 participants (white outline above and in main text Figures 3A and 5A). We note that this outline is indeed larger than an individual participant's actual habenula volume.

Our manual habenula tracing was guided by anatomical landmarks and visible tissue contrast described by Lawson and colleagues (2013). Given that our anatomical scans were of standard resolution (1mm$^3$, as opposed to higher resolution [0.77 mm$^3$] as in Lawson et al., 2013) and that our intent was to provide a visual frame of reference for the functional outcomes regarding the approximate location of the habenula across the entire sample (as opposed to the precise characterization of structural volume as in Lawson et al., 2013), our methodology for tracing the habenula was not as rigorous as that described by Lawson *et al.* (2013). Rather, our manual tracing procedures were guided by their anatomical descriptors. Specifically, we began by identifying the posterior boundary of the habenula by first locating the coronal slice in which the pineal stalk was visible in the third ventricle. Moving forward (rostral) approximately one slice, the posterior portions of the habenula became visible and slightly extended into the third ventricle. The posterior boundary corresponded to slices in which the posterior commissure and habenula were present. The medial and lateral boundaries were distinguished from neighboring cerebral spinal fluid (CSF) and thalamic grey matter, respectively, by the habenular nuclei's brighter tissue contrast associated with greater white matter density. More precisely, when moving rostrally, the medial boundary was defined by the CSF of the third ventricle and the lateral boundary was defined by the tissue contrast (darker voxels) of the thalamic stria medullaris. Finally, the anterior boundary was identified as the most anterior slice where there was no longer brighter habenular tissue visibly extending into the third ventricle CSF and the dorsal aspect of the stria medullaris was visible.

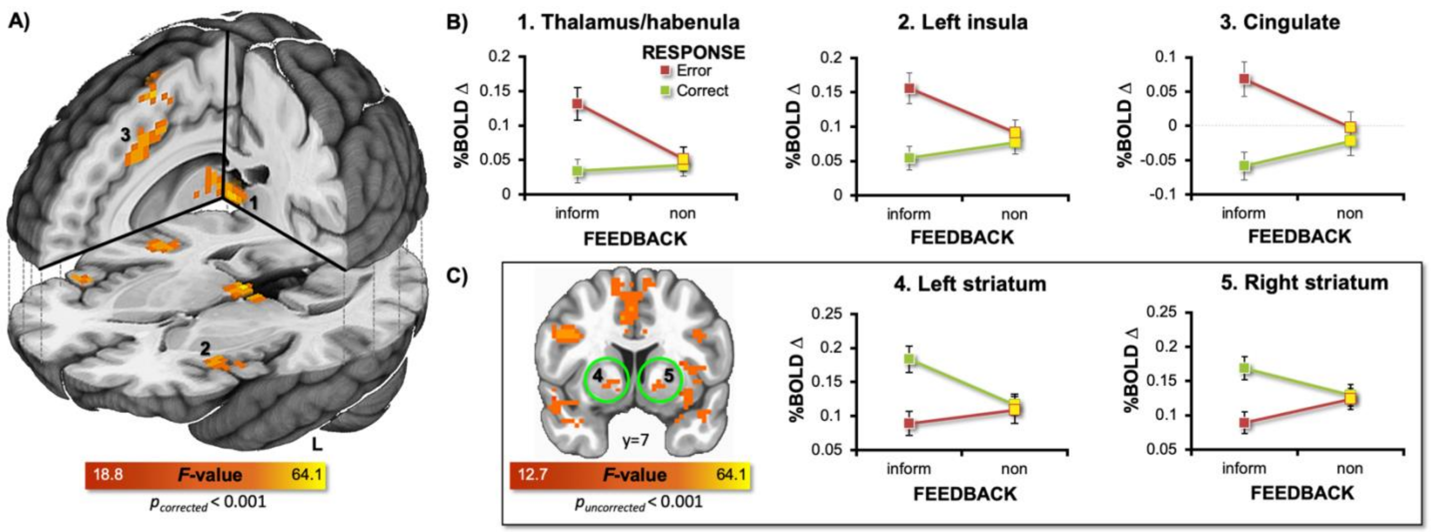


**Figure S2. Task effect: Whole-brain RESPONSE * FEEDBACK interaction analysis.** We performed a whole-brain, 2 (FEEDBACK: informative [i] vs. non-informative [n]) * 2 (RESPONSE: error [E] vs. correct [C]) repeated-measures ANOVA utilizing AFNI's 3dMVM program and the session-averaged $\beta$ coefficients associated with specific task events (iE, iC, nE, nC trials). **(A)** We observed significant RESPONSE x FEEDBACK interaction effects notably in the thalamus encompassing the habenula (94 voxels, X = 2, Y = -23, Z = 5), bilateral anterior insula (left: 142 voxels, X = -38, Y = 18, Z = 7) ; right: 84 voxels, X = 43, Y = 19, Z = 9), and the dorsal cingulate (162 voxels, X = 2, Y = 16, Z = 42) ($p_{corrected} <$ 0.001, $p_{voxel\text{-}wise} < 0.0001$, cluster-extent: 15 voxels). **(B)** Consistent with the whole-brain, one-sample *t*-test assessment reported in the main text (Fig. 3, [iE – iC] contrast values), these interactions in the habenula, insulae, and cingulate clusters were driven by increased activity following negative (informative) feedback (iE > iC), yet no difference between error and correct trials followed by noninformative feedback (nE = nC). **(C)** At an uncorrected threshold ($p_{voxel\text{-}wise} < 0.001$), we observed a RESPONSE * FEEDBACK interaction in the bilateral ventral striatum (left: 6 voxels, X = -16, Y = 8, Z = -4; right: 9 voxels, X = 16, Y = 9, Z = -2). These interactions in the striatum were driven by increased activity following positive (informative) feedback (iE < iC), yet no difference between correct and error trials followed by noninformative feedback (nE = nC). Collectively, these interaction outcomes provided quantitative support isolating the critical task manipulation to the type of feedback (i.e., negative vs. positive) as opposed to the type of response (i.e., error vs. correct trials).

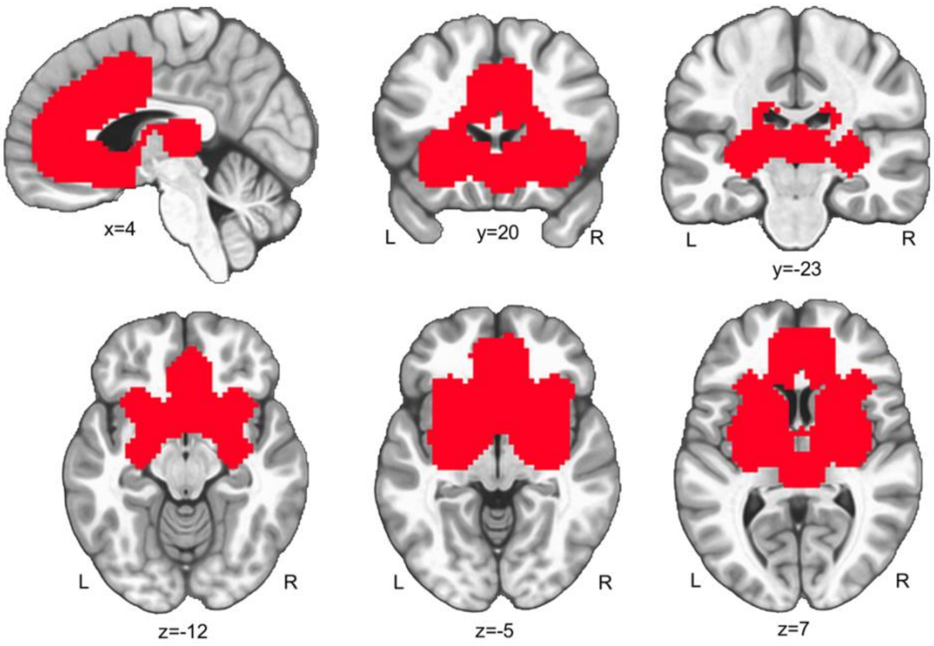


**<u>Figure S3.</u>** ***A priori* composite mask of interest.** Given our *a priori* focus on brain regions previously linked with addiction, nicotine withdrawal, and feedback processing within the mesocroticolimbic circuitry (Fedota et al., 2015; Ullsperger and von Cramon, 2003; Lesage et al., 2017; Sutherland et al., 2012), we utilized a small volume of interest approach when characterizing group and drug effects on brain activity during the motion prediction task. The volume of interest was created by summing the anatomical masks of 16 regions from the Desai maximum probability map (DD_Desai_PM, Destrieux et al., 2010) which included the left and right nucleus accumbens, caudate, putamen, pallidum, anterior insula, anterior cingulate, middle anterior cingulate, and subcallosal cingulate. Additionally, left and right habenula masks (not available in the Desai map) were included and defined by placing a 3mm radius sphere around the left and right habenula coordinates reported in the original implementation of the motion prediction task (Ullsperger and von Cramon, 2003). All 18 regional masks were summed, dilated by three voxels, and then eroded by one voxel to fill any "holes" in the composite mask. Finally, the lateral ventricles and portions of the corpus callosum, that were included in the composite mask after dilation, were subtracted. AFNI's 3dClustSim algorithm (implemented with the spatial autocorrelation function correction) was subsequently used to estimate appropriate cluster-extent corrections to arrive at a family-wise error (FWE) corrected threshold of $\alpha < 0.05$ across the mask. Within the volume of interest (7,684 voxels) and utilizing a voxel-level threshold of $p < 0.001$ with 2-sided thresholding and second nearest neighbor clustering options (NN = 2, https://afni.nimh.nih.gov/pub/dist/doc/program_help/3dClustSim.html), a cluster-extent threshold of 9 voxels yielded a $p_{corrected} < 0.05$ threshold.

For the whole-brain task effect, we utilized a 90% probability group mask of the functional data across all participants and scan sessions (51,074 voxels). Using 3dClustSim and a voxel-level threshold of $p < 0.0001$ with 2-sided thresholding and second nearest neighbor clustering options (NN = 2), a cluster-extent threshold of 15 voxels, yielded a $p_{corrected} < 0.001$ threshold.

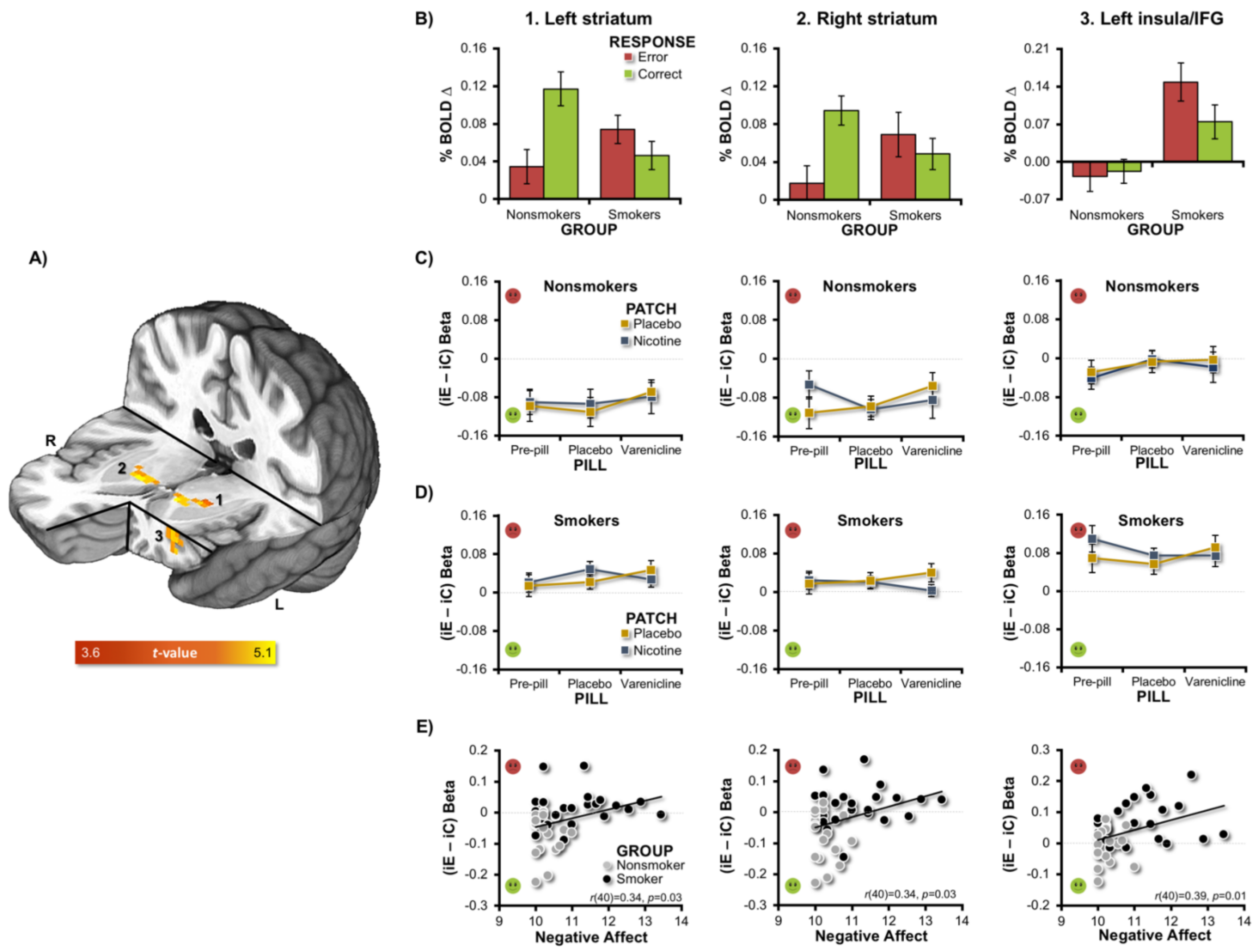


**Figure S4. Group effects: Additional characterization of regions showing smoker versus nonsmoker differences in feedback responsivity.** **(A)** Group differences in [iE – iC] contrast values were observed in the bilateral ventral striatum and left anterior insula/inferior frontal gyrus (IFG). Duplicated from **main text Fig. 4A** for context here. **(B)** To further assess the nature of the group differences observed when considering the [iE – iC] contrast values (i.e., a difference score), we separately plotted the beta weights from iE and iC trials for the nonsmoker and smoker groups. Given a direct statistical comparison of these beta weights would constitute a circular analysis (Kriegeskorte et al., 2009), we offer a qualitative description of these beta weights. When considering the left ventral striatum, the mean difference between smokers and nonsmokers following iC (positive) feedback (green bars) yielded a large effect size (Cohen's $d$ = 0.96) such that BOLD signal change was reduced by 60.3% among smokers. In contrast, the mean difference between smokers and nonsmokers following iE (negative) feedback (red bars) yielded a small-medium effect size (Cohen's $d$=0.39). This qualitative assessment suggested that the group difference observed in the [iE – iC] difference scores was predominately driven by smoker's reduced responsivity to positive feedback. When considering the right ventral striatum, the mean difference between smokers and nonsmokers following iC (positive) feedback (green bars) yielded a medium-large effect size (Cohen's $d$ = 0.64) such that BOLD signal change was reduced by 48.5% among smokers. The mean difference between smokers and nonsmokers following iE (negative) feedback (red bars) yielded a medium effect size (Cohen's $d$=0.54). In sum, when assessing the beta weights separately for iE and iC trials a larger effect size between groups was observed following iC (positive) feedback such that smoker's BOLD signal change was reduced by 60% in the left and 48% in the right ventral striatum. **(C, D)** To assess for state-level drug effects within these clusters, we separately plotted the [iE – iC] contrast values across each drug condition. While we observed a high degree of consistency in the contrast values (speaking to replicability), we did not observe any indication of nicotine or varenicline effects within these clusters among nonsmokers (C) nor smokers (D) ($p$'s > 0.15). **(E)** We conducted additional exploratory analyses across all participants to relate task-related brain activity with self-reported positive and negative affect. Specifically, participants completed the PANAS at each study visit (9 times total: 1 orientation, 6 neuroimaging, 2 neurocognitive visits). Given we were interested in trait-levels of affect (paralleling the session-averaged, trait-level assessments of brain activity), we computed a session-averaged metric of positive and negative affect. We were interested in an individual's general level of affect as opposed to session-to-session fluctuations. Across all participants, reduced positive feedback responsivity (less negative/more positive values) in the left and right ventral striatum was positively correlated with higher trait-levels of negative affect (left: $r[40] = 0.34$, $p = 0.03$; right: $r[40] = 0.34$, $p = 0.03$, given the exploratory nature of these analyses, uncorrected $p$-values are reported). In the left insula/IFG greater negative feedback responsivity (higher positive values) was positively correlated with higher levels of negative affect ($r[40] = 0.39$, $p = 0.01$). These brain-behavior relationships were not observed when considering positive affect (left striatum: $r[40] = -0.29$, $p = 0.07$, right striatum: $r[40] = -0.19$, $p > 0.2$, insula/IFG: $r[40]= -0.12$, $p > 0.4$; data not shown).

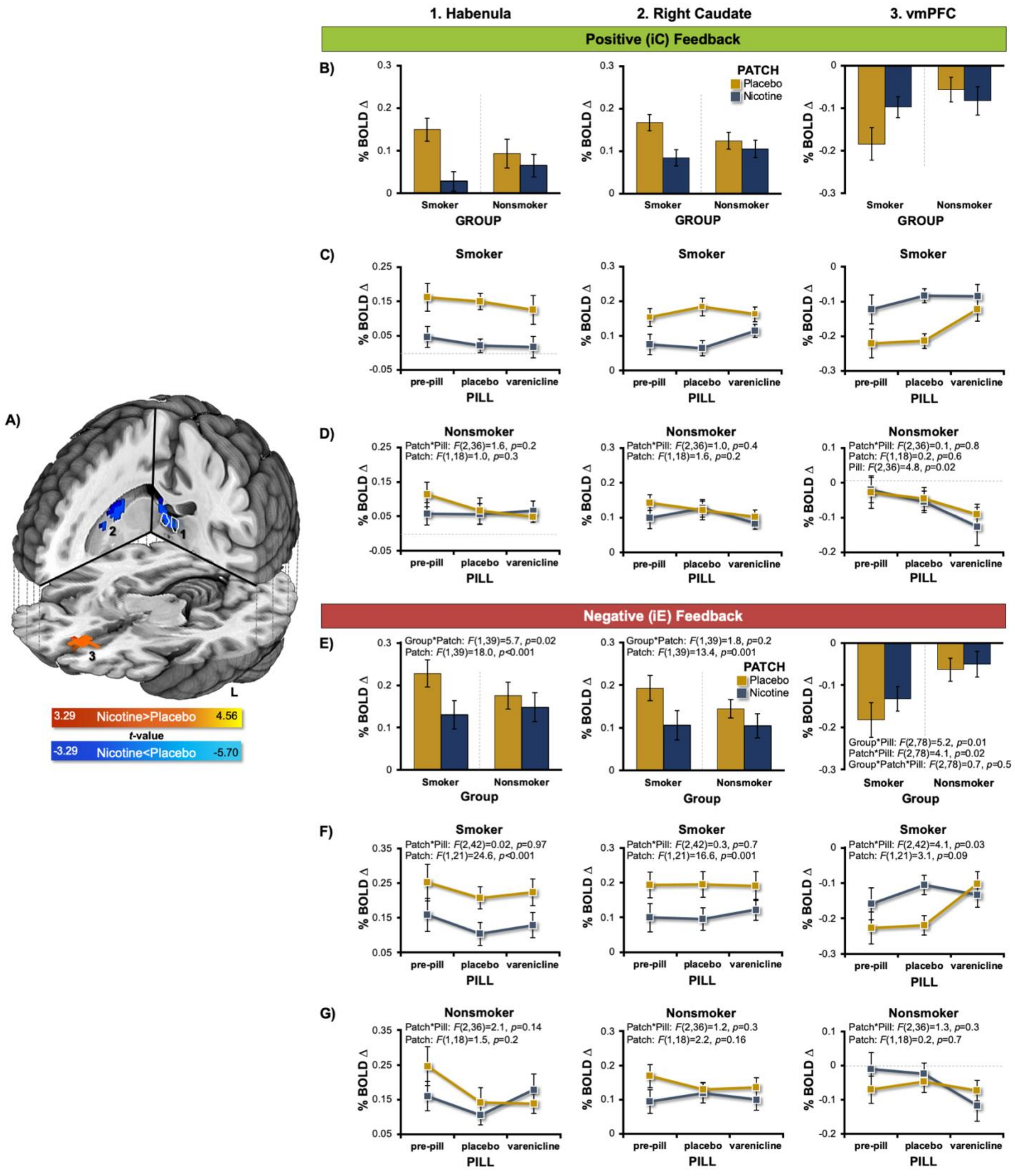


**Figure S5. Drug effect: Pharmacologically-induced alterations following *both* positive (iC) and negative (iE) feedback across all sessions for the two participant groups (smokers, nonsmokers).** To facilitate comprehensive interpretation of pharmacological effects, we considered activity alterations in select ROIs following *both* positive (iC) and negative (iE)

feedback across all sessions and the two participant groups. **(A)** Select ROIs were identified as those showing acute nicotine-induced effects among smokers (AFNI's 3dLME), namely the habenula (outlined in white), left and right caudate, and cingulate gyrus (cold colors) and vmPFC (hot colors) following positive (iC) feedback (duplicated from main text Figure 5A for context here). We then extracted the $\beta$ coefficients from these ROIs separately following *both* positive (iC) and negative (iE) feedback for all 6 sessions/drug conditions and across both smokers and nonsmokers thereby allowing for graphical and/or selective statistical examination. Given that a *selective* statistical analysis of both groups' iE $\beta$ coefficients was unrelated to the statistical test used to select voxels of interest (i.e., the 3dLME of smokers' iC $\beta$ coefficients) (Kriegeskorte et al., 2009), we assessed iE $\beta$ coefficients extracted from identified ROIs in a GROUP * PATCH * PILL mixed-effects ANOVA. For a similar reason (i.e., smokers and nonsmokers were independent samples), nonsmokers' iC $\beta$ coefficients from identified ROIs were assessed in a PATCH * PILL repeated-measures ANOVA. Smokers' iC $\beta$ coefficients (i.e., those used in the statistical test for voxel selection) were extracted only for graphical display.

**Positive (iC) Feedback. (B)** When considering positive feedback, graphical inspection of iC $\beta$ coefficients from select ROIs indicated that nicotine-induced alterations were observed among smokers, but not nonsmokers in the habenula, right caudate, and vmPFC (duplicated from main text Figure 5B for context here). **(C)** Allowing for more comprehensive interpretation, smoker's iC $\beta$ coefficients were then plotted across all 6 sessions. While a selective statistical assessment of smokers' iC $\beta$ coefficients would constituent a circular analysis (Kriegeskorte et al., 2009), visual inspection of the habenular outcomes revealed a consistent nicotine-induced decrease (vs. placebo patch) in habenular activity across all 3 pill conditions (pre-pill, placebo-pill, varenicline-pill). When considering activity in the right caudate and vmPFC, robust nicotine-induced decreases were observed under the pre-pill and placebo-pill conditions, yet an attenuated nicotine versus placebo patch difference was observed under varenicline-pill conditions. These outcomes in the caudate and vmPFC are generally consistent with the known pharmacodynamic actions of nicotine (full agonist) and varenicline (partial agonist) at $\alpha 4\beta 2$ nAChRs (Rollema et al., 2007) and similar to previous observations when assessing heart rate (Sutherland et al., 2013a), behavioral measures (Sutherland et al., 2013a, Lesage et al., 2017, Carroll et al., 2015) including those from the current task (i.e., main text Figure 2C), as well as task-based fMRI (Lesage et al., 2017, Fedota et al., 2015, Sutherland et al., 2013a) and resting-state fMRI outcomes (Sutherland et al., 2013b). **(D)** Selective statistical assessment of nonsmokers' iC $\beta$ coefficients, failed to detect nicotine-induced alterations in the habenula, caudate, or vmPFC. However, a PILL main effect was detected in the vmPFC which was driven by varenicline-induced enhanced deactivations under active pill conditions. Taken together, these outcomes support the interpretation that nicotine-induced alterations following positive feedback were observed among smokers, but not nonsmokers particularly in the habenula.

**Negative (iE) Feedback. (E)** When considering negative feedback, acute nicotine (vs. placebo patch) was similarly associated with reduced activation of the habenula among smokers, but not nonsmokers. This was indicated by a significant GROUP * PATCH interaction ($F[1,39] = 5.7$, $p = 0.02$), which was followed by within-groups repeated-measures ANOVAs. We observed a nicotine-induced reduction of habenula activity among smokers (PATCH main effect: $F[1,21] = 24.6$, $p < 0.001$, **F**), but not nonsmokers (PATCH main effect: $F[1,18] = 1.5$, $p = 0.2$, **G**). For completeness, we also conducted this same series of statistical assessments for the caudate and vmPFC ROIs. With respect to the caudate, although the GROUP * PATCH interaction failed to reach significance ($F[1,39] = 1.8$, $p = 0.2$, **E**), we detected a nicotine effect among smokers ($F[1,21] = 16.6$, $p = 0.001$, **F**), but not nonsmokers ($F[1,18] = 2.2$, $p = 0.16$, **G**). With respect to the vmPFC, we observed a PILL * PATCH interaction among smokers ($F[2,42] = 4.1$, $p = 0.03$, **F**), but not nonsmokers ($F[2,36] = 1.3$, $p = 0.3$, **G**) in the absence of a significant GROUP * PATCH * PILL interaction ($F[2,78] = 0.7$, $p = 0.5$). Taken together, these outcomes were consistent with the interpretation that nicotine-induced alterations following *both positive as well as negative feedback* were observed among smokers, but not nonsmokers particularly in the habenula.

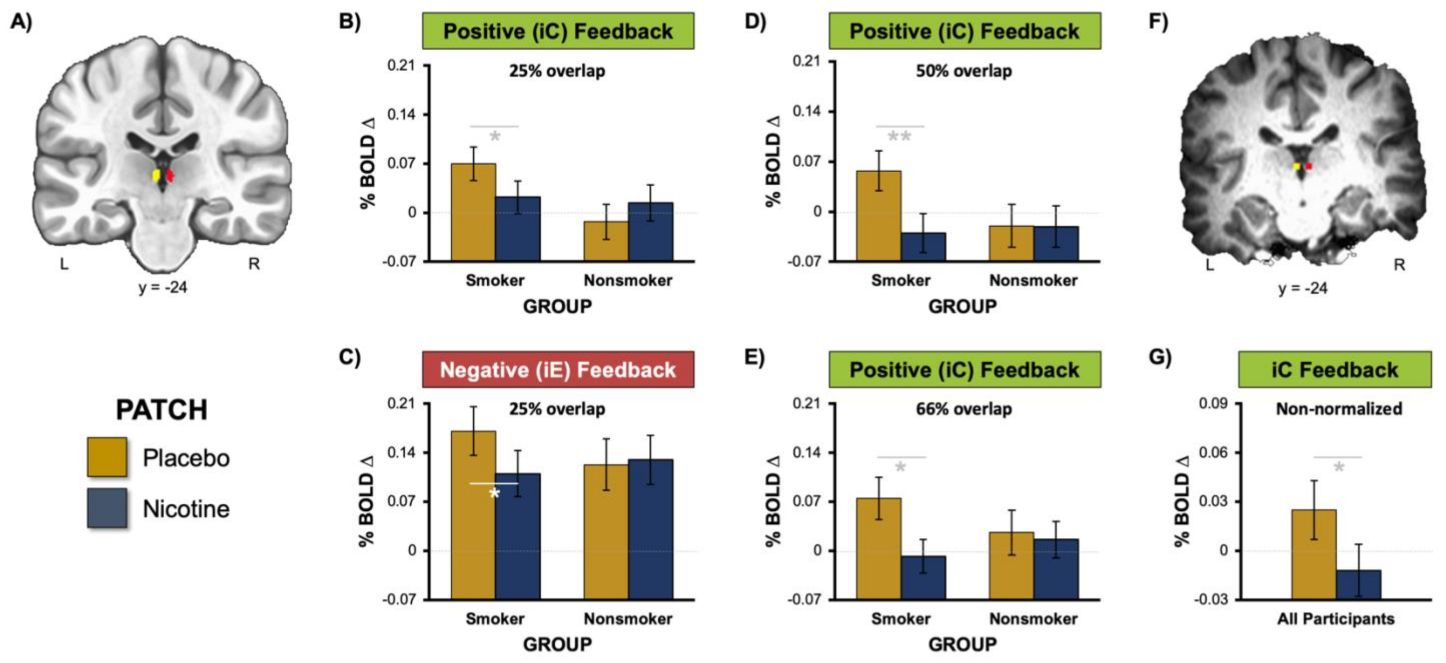


**<u>Figure S6.</u> Drug-effects: ROI-based analyses utilizing anatomically-defined left and right habenula locations to assess nicotine-induced alterations following *both* positive (iC) and negative (iE) feedback.** To provide additional support for the interpretation that nicotine administration reduced habenula activity following positive and negative feedback among smokers (but not nonsmokers), we utilized two different approaches involving ancillary ROI-based analyses. In the first approach, we used the <u>post-normalized fMRI data</u> for assessment and in the second approach we utilized the <u>non-normalized fMRI data</u>. For both approaches, we began by creating ROI masks utilizing each participant's anatomically-defined habenula locations (see: **Supplemental Fig. S1** for details).

In the <u>post-normalized ROI</u> assessments, we then created separate left and right habenula masks defined as voxels where at least 25% of the participants showed overlapping habenula locations (**<u>A</u>**). We acknowledge that ROI-based assessments of the habenula are challenging. On the one hand, given the small size of the habenula (~30mm3, Lawson et al., 2013) and the resolution of our resampled fMRI voxels (27mm3, 3mm isotropic), it would seem desirable to construct ROIs as small as possible to mitigated the influence of partial voluming issues (i.e., the impact of non-habenula signal). On the other hand, given the "noisiness" of the BOLD signal, particularly when considering deep brain voxels at 3T, utilizing a single-voxel ROI would seem to increase susceptibility to inaccurate estimates resulting from noise, regardless of the level of care exercised when selecting that voxel. Typical strategies to deal with such noise (e.g., spatial smoothing, averaging over multiple voxels, spatial normalization to minimize the influence of individual anatomical differences) increase the likelihood of mixing habenula with non-habenula signal. As such, our rationale for selecting the admittedly arbitrary threshold of 25% participant overlap was to seek a balance between these two competing challenges (partial voluming vs. noise reduction). Specifically, a 25% threshold yielded ROI masks composed of 4 functional voxels in the left hemisphere and 4 voxels in the right hemisphere. Average beta weights from voxels within these masks were extracted for both positive (iC) and negative (iE) feedback across all 6 sessions and both participant groups. We then performed separate 4-way mixed-effects ANOVAs in SPSS with factors for GROUP (smoker, nonsmoker), PATCH (nicotine, placebo), PILL (varenicline, placebo, pre-pill), and HEMISPHERE (left, right) for the two feedback types.

When considering positive feedback **<u>(B)</u>**, the only effect to emerge as significant in the omnibus analysis was a GROUP * PATCH interaction ($F[1, 39] = 6.3$, $p = 0.02$). Given that there were no significant main or interaction effects with the HEMISPHERE factor (or any other factor, $p$'s > 0.15), we collapsed across the left and right hemispheres when performing *post hoc* follow-up assessments. These *post hoc* assessments indicated that the GROUP * PATCH interaction was driven by a nicotine-induced reduction in habenula activity among smokers ($t[21] = -2.1$, $p = 0.046$), but not among nonsmokers ($t[18] = 1.6$, $p = 0.14$). When considering negative feedback **<u>(C)</u>**, we also observed a significant GROUP * PATCH interaction ($F[1, 39] = 4.6$, $p = 0.04$). *Post hoc* assessments again indicated that the GROUP * PATCH interaction was driven by a nicotine-induced reduction in habenula activity among smokers ($t[21] = -3.1$, $p = 0.005$), but not nonsmokers ($t[18] = 0.3$, $p = 0.8$).

To increase confidence in the 25%-overlap ROI analyses, we then considered estimates of habenula activity following positive (iC) feedback from the post-normalized fMRI data at different participant overlap thresholds (i.e., 50% and 66%). Increased thresholds lead to smaller ROIs; a 50% threshold yielded ROIs composed of 3 left and 1 right hemisphere voxel and a 66% threshold yielded a ROI composed of 1 left hemisphere voxel (no voxels in the right hemisphere reached this threshold). We again performed a 4-way mixed-effects ANOVA with factors for GROUP, PATCH, PILL, and HEMISPHERE for the 50%-threshold assessment and dropped the HEMISPHERE factor for the 66%-threshold assessment. When considering the 50% threshold **(D)**, we observed an even more robust GROUP * PATCH interaction ($F[1, 39] = 15.5$, $p < 0.001$) again driven by a nicotine-induced reduction in habenula activity among smokers ($t[21] = -6.1$, $p < 0.001$), but not nonsmokers ($t[18] = -0.04$, $p = 0.97$). When considering the 66%-threshold **(E)** we also observed a GROUP * PATCH interaction ($F[1, 39] = 6.9$, $p = 0.012$) driven by a nicotine-induced reduction in habenula activity among smokers ($t[21] = -4.7$, $p < 0.001$), but not nonsmokers ($t[18] = -0.5$, $p = 0.65$). Taken together, when utilizing increasing smaller ROIs for the assessment of habenula activity, we observed similar outcomes leading to the same interpretations reported in the main text Figure 5. Specifically, these outcomes indicated that nicotine administration reduced habenula activity following performance feedback among smokers, but not nonsmokers.

In the non-normalized ROI assessments, we created ROIs by identifying the center of mass coordinates separately for the left and right hemispheres from the habenula mask manually drawn on each participants' T1-weighted structural image. These coordinates were then used to select the single voxel from the resampled, co-registered and non-normalized fMRI corresponding to the habenula. **(F)** The single left and right voxel corresponding to the habenulae-designated functional voxels from one example participant. **(G)** In a 4-way, GROUP * PATCH * PILL * HEMISPHERE ANOVA, we observed a slightly different pattern of outcomes than that from the post-normalized assessments. Specifically, we did not detect a significant GROUP x PATCH interaction ($F[1, 39] = 0.4$, $p = 0.55$), but rather a significant PATCH main effect ($F[1, 39] = 6.7$, $p = 0.014$). Stated differently, in this assessment, nicotine-induced decreases in habenular activity were similar among smokers and nonsmokers. We speculate that these outcomes may be influenced by noise when assessing non-normalized fMRI data. Nonetheless, these outcomes are consistent with the more general interpretation that acute nicotine administration is capable of down-regulating habenula activity.

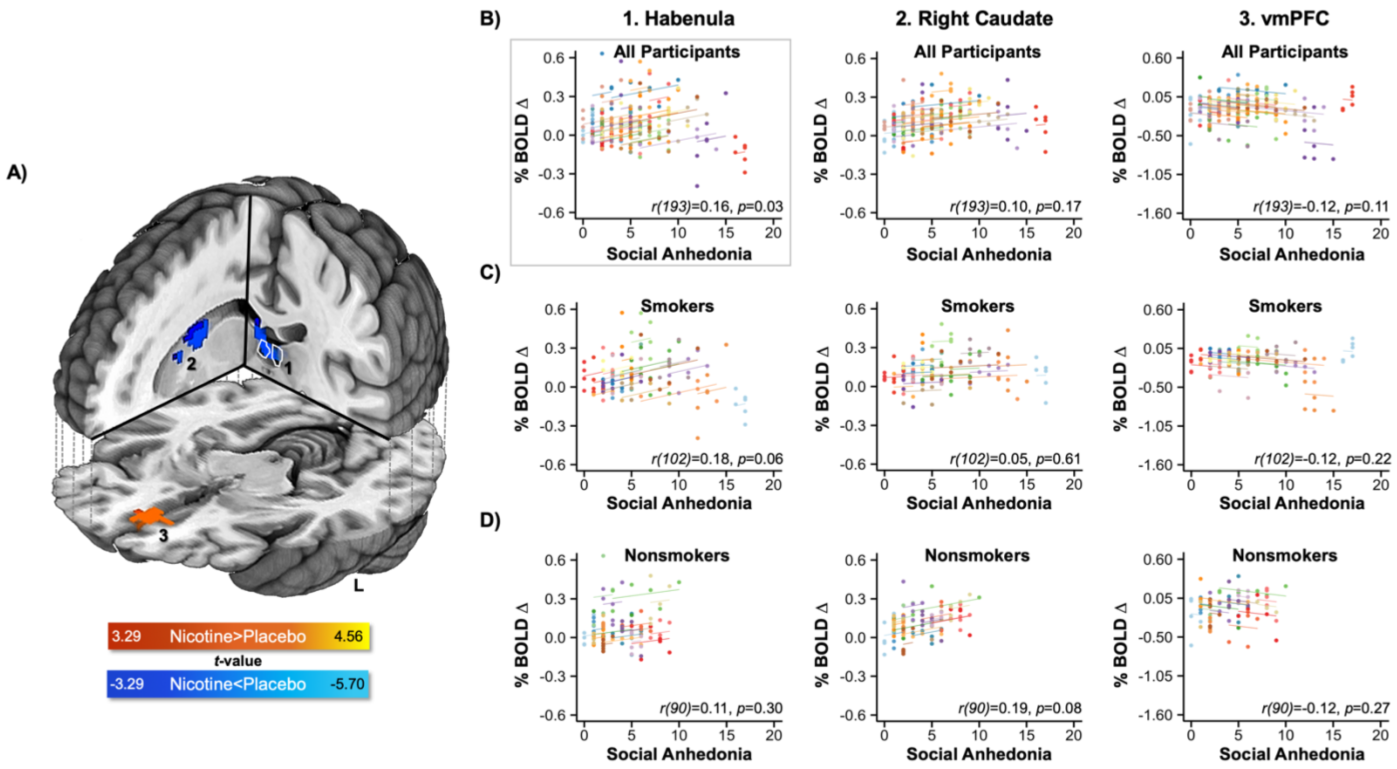


**Figure S7. Drug effects: Additional characterization of regions showing nicotine-induced alterations.** (A) Acute nicotine administration to smokers was associated with reduced activation in the habenula (outlined in white), left and right caudate, and cingulate gyrus (cold colors) as well as reduced deactivation of the ventral medial prefrontal cortex (vmPFC) on iC trials (hot colors). Duplicated from main text Figure 5A for context here. **(B-D)** We conducted exploratory repeated measures correlation (RMcorr) analyses to link task-related brain activity with self-reported state-levels of social anhedonia. Specifically, participants completed the Revised Social Anhedonia Scale during each of the six neuroimaging visits. Given we were interested in state-levels of anhedonia (i.e., session-to-session fluctuations), we used RMcorr to assess the within-individual relationship between brain activity following positive feedback and social anhedonia self-reported each session by all participants (B), among smokers only (C), and among nonsmokers only (D). When considering all participants, the RMcorr indicated that as an individual participant's habenula activity increased across sessions, so too did their self-reported level of social anhedonia ($p$ = 0.03, given the exploratory nature of these analysis uncorrected $p$-values are reported). Indicative of regional specificity, a similar within-individual relationship was not observed when considering the right caudate ($p$ = 0.17) or vmPFC activity ($p$ = 0.11). When considering cohorts of participants separately (C, D), no significant associations were detected.
